# The multi-scale geometry of neural encoding

**DOI:** 10.64898/2026.10.22.761206

**Authors:** Simone Azeglio, Steeve Laquitaine, Arianna di Bernardo, Ulisse Ferrari, Matthew Chalk

**Affiliations:** Institut de la Vision, Sorbonne Université, Paris, France; Laboratoire des Systèmes Perceptifs, École Normale Supérieure - PSL, Paris, France; Laboratoire de Neurosciences Cognitives et Computationnelles, École Normale Supérieure - PSL, Paris, France; Centre National de la Recherche Scientifique (CNRS); Institut national de la santé et de la recherche médicale (INSERM)

**Keywords:** neural coding, mutual information, representational geometry, vision, population coding, efficient coding, Fisher information, diffusion models

## Abstract

As recorded neural populations become larger and artificial networks more powerful, there is growing interest in understanding and comparing what different populations encode. Representational geometry offers a powerful way to characterise this: it remaps stimulus space to stretch well-encoded stimulus differences while shrinking poorly encoded ones. However, this remapping can be constructed in many ways from neural responses. Here, we introduce a multi-scale representational geometry, derived from first principles, that captures how neurons encode stimulus differences at every scale. It is exactly related to mutual information, so independent of decoding assumptions. The more a stimulus difference contributes to information, the more the geometry stretches it. Applied to spatial, motor and visual codes, it shows how neurons combine to encode information, how areas differ, and which features are best encoded. A single framework thus maps the structure of encoded information, uniting geometric and information-theoretic views of neural coding.

## INTRODUCTION

Our ability to perceive the world depends on how neurons in our brain respond to sensory stimuli. Stimuli that reliably evoke different neural responses can be distinguished; those that do not cannot. Recent years have seen massive growth in our ability to record from large populations of neurons and measure how they respond to different stimuli^1–4^. Despite this, characterising what neural populations tell us about the world — what stimulus features are encoded and which are not — remains a challenge.

Representational geometry offers a way forward by remapping stimulus space according to the neural code: stimuli that evoke distinct responses are pulled apart, while those that evoke similar responses are drawn together^5,6^. Stimulus differences that the code preserves are thus stretched, and those it discards are compressed (Fig. 1A). This allows neural codes to be compared directly: across brain areas^7–10^, between biological and artificial systems^11–14^, with perception^15–17^, and over the course of learning and adaptation^9,18,19^.

**Figure 1.**
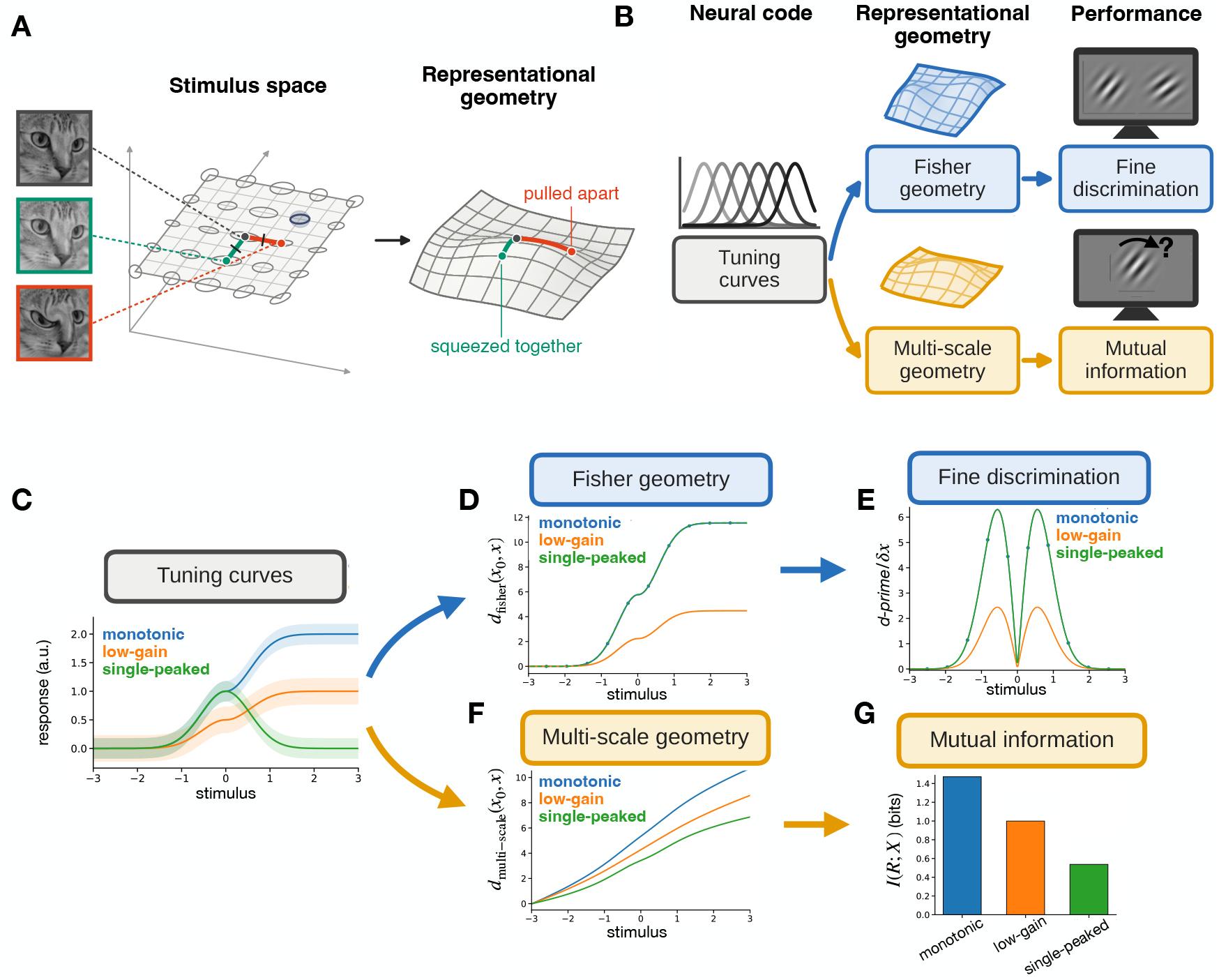
Multi-scale representational geometry. (A)Left: two images (red, green) differing equally from a reference image (black) in their pixels (mean-squared difference). At every stimulus, a metric tensor (ellipses) describes how well neurons encode each stimulus dimension. The elongated ellipse at the reference image magnifies distances towards the red image and shrinks distances towards the green image, indicating that neurons encode the first dimension better than the second. Right: the resulting representational geometry, in which distances reflect how well stimuli are distinguished by the neural code. Here, the red image lies far from the reference and the green image close to it, even though both perturbations are equi-distant in pixel space. (B) A neural code induces a geometry over stimulus space. The Fisher geometry predicts fine discrimination performance, e.g. in discriminating two nearby orientations. In the multi-scale geometry, distances are exactly related to the mutual information between neural responses and stimuli e.g. determining how well orientation can be estimated. (C) Three example tuning curves. Solid lines show mean responses and shading is variability. (D) Fisher distances to a stimulus at *x*0 = −3 for each tuning curve. Monotonic and single-peaked tuning have identical Fisher geometries so their curves are overlapping. (E) Fine discrimination performance (normalized d′), predicted by the Fisher information, is also identical for the monotonic and single-peaked tuning curves. (F) Distances in the multi-scale geometry are reduced for the single-peaked tuning curve relative to the monotonic curves because it encodes the stimulus ambiguously. (G) Mutual information shows the same ordering as the multi-scale geometry, with monotonic tuning curves conveying more information than the single-peaked tuning curve.

However, there is more than one way to define distances between stimuli from the neural responses they evoke. This choice matters: each definition yields a different geometry, and so a different account of what a population encodes. One possibility is to define distances based on the mean activity evoked by different stimuli^20–23^. Yet, this ignores response variability, which strongly affects what a population can encode^24–26^. Another is to measure how accurately a particular decoder can tell stimuli apart, as with linear classifiers or the cross-validated Mahalanobis distance used in representational similarity analysis^7,27,28^, but this ties the geometry to an assumed readout^29,30^. Information geometry avoids both problems by constructing distances from Fisher information, which measures how reliably nearby stimuli can be discriminated from the full distributions of neural responses^31–34^. But, as we show, a geometry constructed this way is blind to whether widely separated stimuli can be told apart: two populations can have identical Fisher information everywhere, yet tell us very different things about the world^35,36^.

Here we introduce a *multi-scale* representational geometry, which captures how well a population encodes stimulus differences across scales, from the finest discriminations to coarse global distinctions. It emerges uniquely from considering how distances between stimuli change for an observer who views them at progressively coarser resolution^37^. To our knowledge, it is the first representational geometry whose distances are exactly related to mutual information: the more a stimulus feature contributes to the information a population encodes, the more the geometry stretches it; where information is lost, stimuli collapse together. Because mutual information depends both on how neurons respond to stimuli and on how often each stimulus occurs, so does the geometry. Stimulus features that rarely vary contribute little information, and are compressed. This links representational geometry directly to normative theories such as efficient coding, which propose that neural codes maximise the information they transmit about the world^38–41^.

Because its distances are tied to mutual information, the multi-scale geometry makes the structure of encoded information visible: how neurons combine to encode information as a population, and how encoded information varies between areas. We demonstrate this in three settings. In entorhinal cortex, the geometry captures how grid cells combine to encode an animal’s position. In motor cortex, it distinguishes two areas by the information they carry about future movement trajectories. Finally, using diffusion models, we extend the framework to high-dimensional natural images, where the geometry identifies which image changes V1 and V4 populations encode best, revealing differences between the two areas. In each case, the Fisher geometry misses this structure, since it reflects only how well small stimulus changes can be discriminated, ignoring both how responses to distant stimuli overlap and how often each stimulus occurs. Together, these results show how a single geometric framework reveals the structure of encoded information across spatial, motor and visual population codes.

## RESULTS

To construct a representational geometry, we must specify how to measure distances between stimuli, ***x*** and ***x***′. The simplest choice, the Euclidean distancem 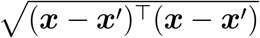, treats all stimulus dimensions equally. A neural population, however, may encode some stimulus changes well and others poorly. To capture this geometrically, we define the distance between two nearby stimuli, ***x*** and ***x*** + ***dx***, as

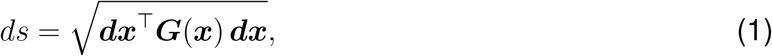

where the matrix ***G***(***x***), called the metric tensor, specifies how distance is measured at each point. Formally, this defines a Riemannian geometry. Setting ***G***(***x***) = ***I*** recovers the Euclidean distance. Our aim, though, is to construct ***G***(***x***) so that it stretches stimulus space along the dimensions that the population encodes well, and compresses it along those it encodes poorly (Fig. 1A). Distances between stimuli, *d*_***G***_(***x, x***′), are then obtained by integrating *ds* along the shortest path connecting them. Thus, if we know the metric tensor ***G***(***x***), we know everything about how space is distorted to form the representational geometry.

Since neural responses, ***r***, are noisy, two stimuli can only be distinguished to the extent that they evoke different *distributions* of responses, *p*(***r*** | ***x***). Distances should therefore reflect how this distribution changes with the stimulus. Information geometry addresses precisely this problem, and identifies the Fisher information matrix as the canonical metric tensor^32^,

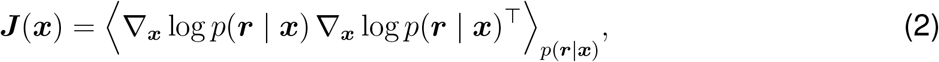

where ⟨·⟩_*p*_ denotes an average over the distribution *p*. The Fisher information quantifies how sensitive the response distribution is to small changes in the stimulus. With ***G***(***x***) = ***J*** (***x***), the distance between two nearby stimuli corresponds to how well an ideal observer could discriminate them from the neural responses (the discriminability, *d*′), which sets a fundamental limit on fine perceptual discrimination^42^. Thus, the Fisher geometry provides a direct link from the neural population code, through the representational geometry, to fine discrimination performance (Fig. 1B).

To illustrate, we consider a single neuron encoding a one-dimensional stimulus *x*, with mean response *f* (*x*) (its tuning curve) and additive Gaussian noise. Here, the Fisher information is proportional to the squared slope of the tuning curve, *J* (*x*) ∝ *f* ′(*x*)^2^. We compare a single-peaked tuning curve with a monotonic curve constructed to have the same slope magnitude everywhere, together with a monotonic curve with reduced gain (Fig. 1C). The single-peaked and matched monotonic curves therefore have identical Fisher geometries (Fig. 1D). Accordingly, fine discrimination performance, *d*′, is identical for both neurons (Fig. 1E). As expected, reducing the gain shrinks distances and lowers *d*′ (Fig. 1D,E).

Yet the single-peaked and matched monotonic neurons convey different amounts of mutual information about the stimulus (i.e. how much, on average, the response reduces uncertainty about it). The single-peaked neuron responds identically to stimuli placed symmetrically on either side of its peak, and so cannot tell them apart; it therefore encodes less information than the monotonic neuron (Fig. 1G). This difference is invisible to the Fisher geometry, and rightly so as far as fine discrimination is concerned: nearby stimuli are discriminated equally well by both neurons. What differs is how well they distinguish widely separated stimuli, which a geometry built from local sensitivity alone cannot capture.

### A multi-scale geometry links neural sensitivity to mutual information

Unlike the Fisher information, the mutual information reflects how well neurons encode stimulus differences at all scales, not just fine perturbations. We sought a representational geometry that does the same. To build it, we imagine an observer who sees the stimulus only at limited resolution, through a blurred version, ***x***_*t*_. This is drawn from a Gaussian distribution centred on the true stimulus, *p*(***x***_*t*_ | ***x***) = *N* (***x***, *t* ***I***), where *t* sets the scale of the blur. Many stimuli, ***x***, are consistent with a given blurred stimulus, ***x***_*t*_. The response distribution that this observer sees is therefore an average of the responses to each stimulus, ***x***, weighted by its probability given ***x***_*t*_,

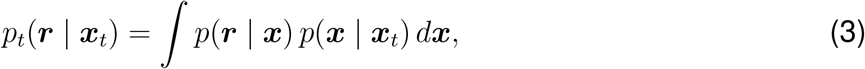

where *p*(***x*** | ***x***_*t*_) ∝ *p*(***x***_*t*_ | ***x***) *p*(***x***), by Bayes’ rule, and *p*(***x***) is the prior distribution of stimuli. The Fisher information of this distribution, ***J*** _*t*_(***x***_*t*_), is the sensitivity that the observer, who only sees ***r*** and ***x***_*t*_, would measure. At fine resolution, it matches the standard Fisher information. As the blur increases, fine stimulus details are lost, and ***J*** _*t*_ comes to reflect how well neurons distinguish coarser stimulus differences. Accumulating this sensitivity across all scales gives,

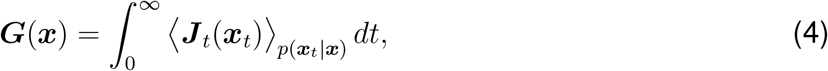

which we call the *multi-scale Fisher information matrix*. We previously showed that this metric tensor defines the unique geometry satisfying a small set of general requirements on how distances should contract as the stimulus is blurred^37^.

The multi-scale geometry is exactly related to the mutual information between stimuli and responses, *I*(*R*; *X*), through the identity^43^

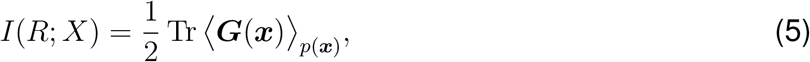

where Tr denotes the trace (Supplementary note). This has a simple geometric interpretation. For two stimuli, ***x*** and ***x***′, the geometry stretches the distance between them by a magnification factor, *m* = *d*_***G***_(***x, x***′)*/*|***x*** − ***x***′|. In the limit of dense sampling from the stimulus prior, the mean squared magnification between neighboring samples is then proportional to the mutual information, *I*(*R*; *X*) ∝ ⟨ *m*^2^ ⟩. The more information the population encodes, the more stimuli are stretched apart; where information is lost, they are squeezed together.

Because it integrates neural sensitivity over all stimulus scales, the multi-scale geometry reflects both the local and global structure of the neural code. Returning to our earlier example, multi-scale distances are reduced for the low-gain curve relative to the matched monotonic one, as for the Fisher geometry (Fig. 1F). Unlike the Fisher geometry, however, they are also reduced for the single-peaked neuron, which encodes stimuli on either side of its peak ambiguously. Across the three tuning curves, distances in the multi-scale geometry follow the same ordering as mutual information (Fig. 1G). Thus, just as the Fisher geometry links the neural code to fine discrimination, the multi-scale geometry links it to the mutual information between stimuli and responses (Fig. 1B).

### The multi-scale geometry reveals how neurons combine to encode information

The information conveyed by a neural population depends both on the sensitivity of individual neurons and on how they combine: whether different neurons carry redundant or complementary information about the stimulus^44^. The Fisher geometry is blind to this. For neurons with independent noise, the population Fisher information is simply the sum over neurons,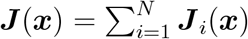.

The Fisher geometry of a population is thus obtained by adding together the contributions of individual neurons, regardless of how their responses relate to one another. Mutual information does not add across neurons in this way, but depends on the pattern of activity across the whole population. Redundant information encoded by different neurons is counted only once. Because the multi-scale geometry is exactly related to mutual information, it inherits this property, revealing how neurons combine to encode the stimulus.

To illustrate, we considered model neurons encoding the orientation and contrast of a visual grating (Fig. 2A–B), where positive and negative contrasts correspond to gratings of opposite polarity. A phase-invariant ‘complex cell’ responds equally to both polarities (Fig. 2C)^45^. Suppose, now, that we split its response between a pair of ‘simple cells’, one responding only to positive contrasts and the other only to negative contrasts (Fig. 2D). At every stimulus, only one of the simple cells responds, and its tuning curve has the same gradient as that of the complex cell, while the silent cell contributes nothing. The summed Fisher information of the pair is therefore identical to that of the complex cell. Yet the pair conveys more information: which of the two cells fires reveals the polarity of the grating, which the complex cell cannot signal.

**Figure 2:**
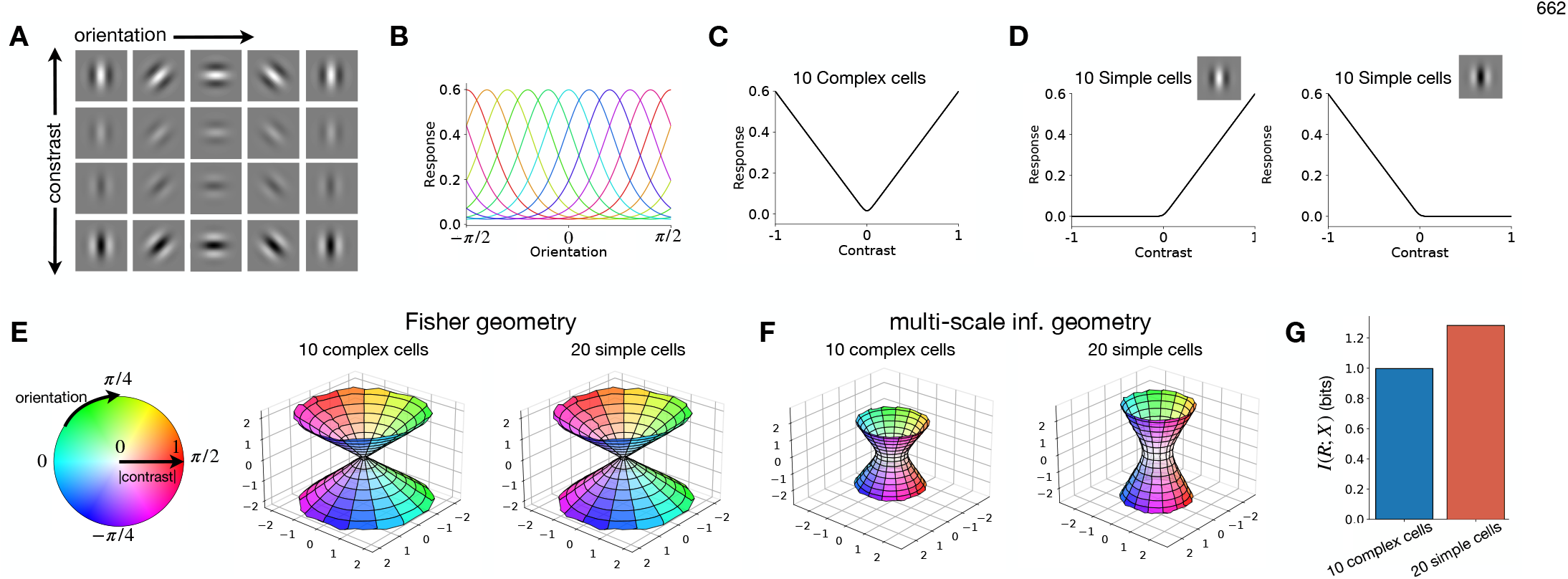
Illustrative model of population coding in primary visual cortex. (A) We consider how neurons in primary visual cortex encode varying orientation and contrast of a visual grating, as shown here. (B) Each curve is the mean response of one neuron versus orientation, at fixed contrast. (C)Population of 10 complex cells whose responses increase equally for positive and negative contrast because they are phase invariant. (D)Population of 20 simple cells: 10 prefer positive contrast and 10 prefer negative contrast. (E)Representational geometry for each population, with stimuli colored by orientation and contrast (left). The Fisher geometry does not distinguish simple from complex cell populations because the summed Fisher information is identical everywhere. (F)By contrast, the multi-scale geometry is expanded between stimuli of opposite contrast for the simple-cell population, reflecting its unambiguous encoding of contrast polarity. (G)This difference is also reflected in the mutual information, which is higher for the simple than for complex-cells.

We compared two populations built in this way: 10 complex cells with different preferred orientations (Fig. 2B–C), and the 20 simple cells obtained by splitting each into a pair (Fig. 2D). In the Fisher geometry, orientations are arranged in a circle whose radius grows with contrast magnitude. The geometry is identical for both populations, since their combined Fisher information is the same (Fig. 2E). In contrast, while the multi-scale geometry has a similar overall shape, it distinguishes the two populations. Stimuli of opposite polarity lie further apart for the simple cells than for the complex cells (Fig. 2F), reflecting the greater mutual information conveyed by the simple cells (Fig. 2G). Moreover, unlike the mutual information, which is a single number, the multi-scale geometry shows where the extra information comes from, since distances between stimuli of opposite polarity are expanded, while distances between orientations are unchanged. We next asked whether the same principle applies in recorded populations, where tuning curves are irregular and heterogeneous across cells. We focussed on grid cells in rat entorhinal cortex^46^. Each grid cell fires at multiple locations arranged in a periodic lattice. A single cell therefore signals the animal’s position ambiguously, since the same response can arise at many different positions. Grid cells are organised into modules: cells within a module share the same grid spacing, while spacing differs between modules. Previous work showed that combining modules with different spacings resolves this ambiguity, allowing the population to specify a unique position over a large range^47^. Grid cells thus provide a test case in which the way neurons combine is well understood. Using recordings from Gardner et al.^48^, we compared two equal-sized populations of cells: a mixed-spacing population, drawn from several modules (Fig. 3A), and a fixed-spacing population, taken from a single module with fine spacing (Fig. 3B).

**Figure 3:**
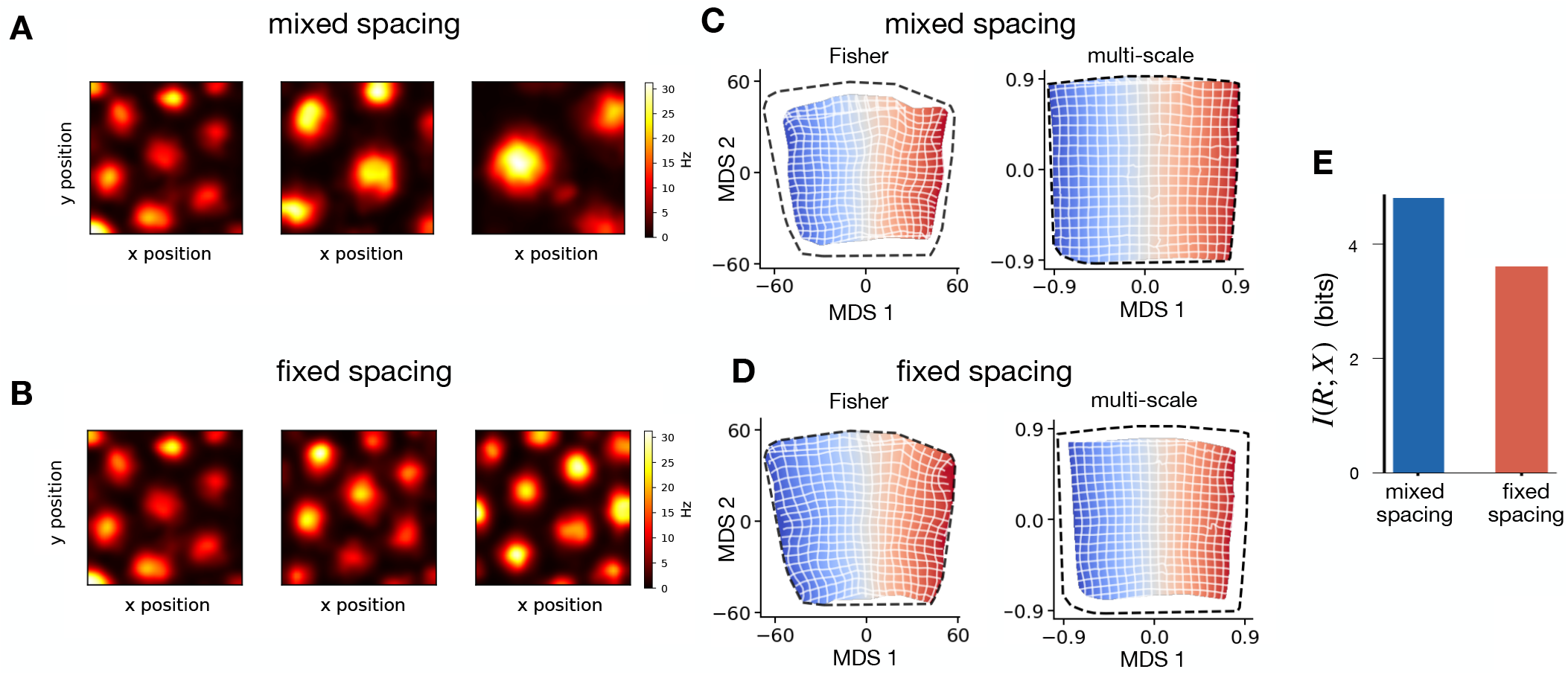
Population coding by grid cells in entorhinal cortex. (A)Three example firing-rate maps, as a function of the animal’s position in a 2D arena, from the mixed-spacing population of 20 grid cells, one from each of the three grid modules, which differ in grid spacing. Data from Gardner et al. ^48^. (B)Three example firing-rate maps from the fixed-spacing population of 20 cells taken from a single module with fixed grid spacing. (C)Representational geometry of the mixed-spacing population, visualized by multidimensional scaling. Left: Fisher geometry. Right: multi-scale geometry. (D)Same as in (C), for the fixed-spacing population. The Fisher geometry occupies a larger area than for the mixed-spacing population, because cells in this module have steeper tuning curves and therefore greater local sensitivity. The multi-scale geometry shows the opposite pattern: it occupies a smaller area than for the mixed-spacing population, where cells from different modules combine to resolve each other’s positional ambiguity. (E)This difference is also reflected in the mutual information, which is higher for the mixed-spacing than for the fixed-spacing population.

In the Fisher geometry, the fixed-spacing population expands the environment more than the mixed-spacing population, so that the same range of positions covers a larger area (Fig. 3C,D, left). This is expected, since fine grid spacing makes individual cells more sensitive to small changes in position, and the Fisher information simply sums these contributions across cells. In the multi-scale geometry, the picture is reversed, with the mixed-spacing population expanding the environment more (Fig. 3C,D, right), reflecting how cells from different modules together pin down the animal’s position, which no single module can do alone. Mutual information is likewise higher for the mixed-spacing population (Fig. 3E). Thus, the multi-scale geometry, but not the Fisher geometry, correctly captures how grid modules combine to encode position.

### The multi-scale geometry reveals differences between motor areas

The same approach can be applied beyond sensory and spatial coding, to how neural populations encode movement. To illustrate this, we analysed recordings from primary motor cortex (M1) and the supplementary motor area (SMA) of macaques turning a hand pedal through repeated cycles (Fig. 4A). Russo et al. found differences in the structure of population activity between the two areas, with SMA tracking aspects of the movement, such as progress through it, that M1 did not^8^. We asked whether such differences would be reflected in the representational geometry of each area, that is, in the distances it induces between movement trajectories.

**Figure 4:**
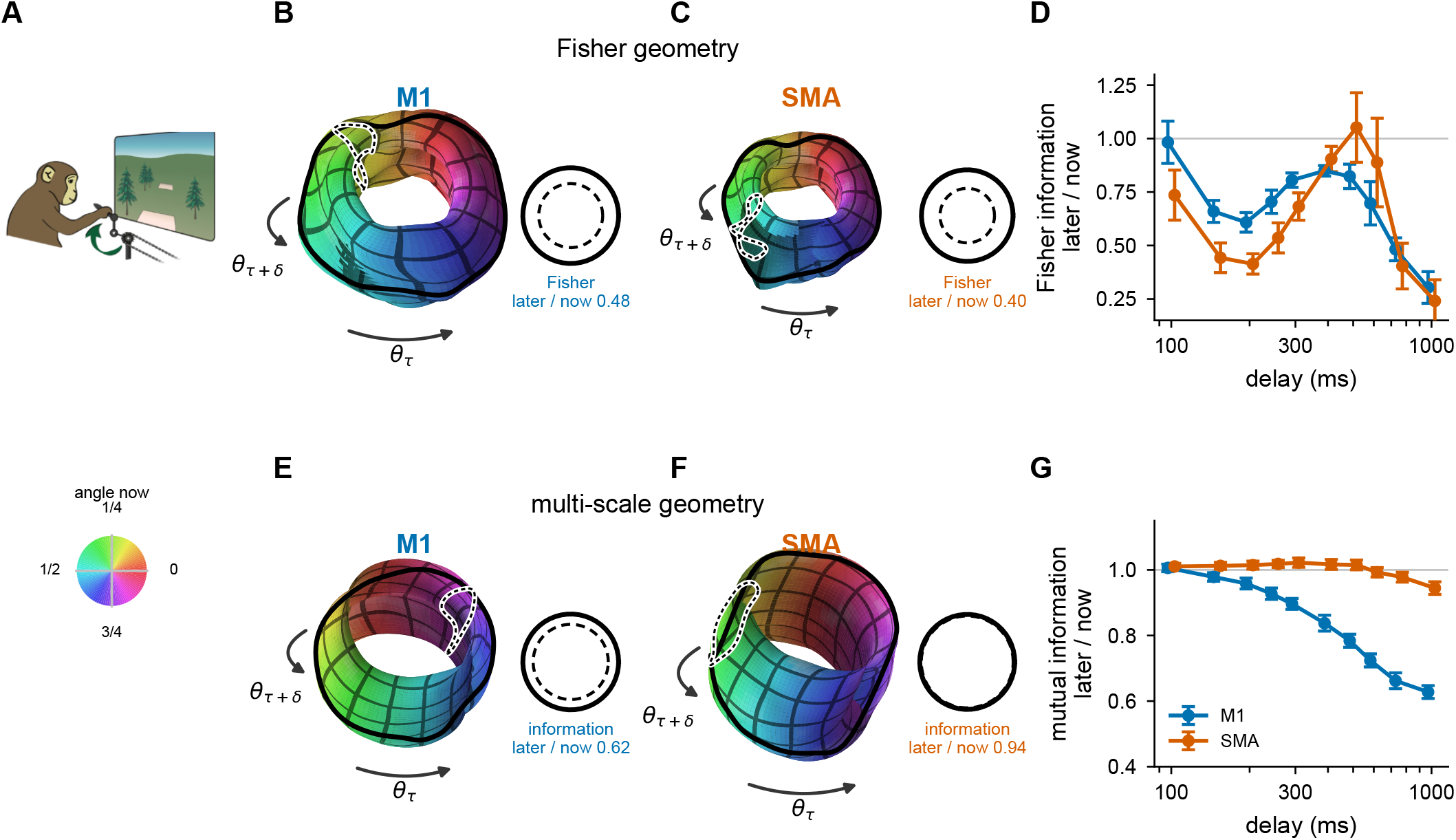
Multi-scale geometry of motor commands, across areas. (A)We considered an experiment where monkeys turned a hand pedal forward or backward through 0.5 to 7 cycles while activity was recorded in primary motor area (M1) and supplementary motor area (SMA). Data from Russo et al. ^8^. (B)We considered a reduced 2-dimensional description of the movement trajectory: its angle at time *τ*, and its angle 750 ms later. The panel show the Fisher geometry of M1, obtained with an MDS projection as in Figure 3. The geometry, pooled over two monkeys, forms a torus, with the current angle of the pedal progressing around the ring, while its angle 750 ms later goes around the tube. Right: circles whose circumferences are set by the Fisher information along each direction of the torus (solid: current angle; dashed: future angle; Methods). (C)Same as (B), for SMA. The overall size of the torus is smaller than for M1. However, the relative size of the two torus diameters remains similar to M1 (right; Fisher information about the future angle relative to the current angle, 0.40 for SMA and 0.48 for M1). (D)Ratio of the Fisher information about the future angle to that about the current angle, against the prediction horizon. Mean of the two monkeys ±s.e.m. (bootstrap over movements). The ratio varies with prediction horizon in a similar way for M1 and SMA, and does not differ consistently between them (statistics for each monkey in Results and Methods). (E–F) Same as (B–C), for the multi-scale geometry. Unlike for the Fisher information, the shape of the torus is different for SMA versus M1: the tube is thicker relative to the overall diameter, implying that SMA encodes more information about the future position, relative to M1 (information about the future and the current angle: 2.35 and 2.51 nats for SMA, 1.57 and 2.55 nats for M1). (G) The mutual information about the future pedal position, divided by the information about its current position, against the prediction horizon. Mean of the two monkeys ±s.e.m. (leave-one-movement-out jackknife). SMA encodes relatively more information about the future, in each monkey, consistent with the multi-scale geometry shown in panels (E–F).

To do so simply, we considered a two-dimensional description of the movement: the position of the pedal at each time, and its position a fixed interval later (the prediction horizon, here 750 ms). Since both variables are angles, this space is a torus, with one circle for current position and one for future position. The circumference of the future-position circle relative to that of the current-position circle shows how well a population encodes upcoming versus current position.

In the Fisher geometry, the tori had similar thickness for the two areas (Fig. 4B-C). The Fisher information about future position, relative to that about current position, varied with prediction horizon in a similar way in both areas, and did not differ consistently between them (Fig. 4D). In one monkey the two areas did not differ (*p* ≥ 0.2); in the other the difference changed sign with horizon (SMA lower up to 300 ms, higher at 400–600 ms and lower again beyond; *p <* 0.001; repeated-measures ANOVA).

In contrast, the multi-scale geometry separated the two areas: the torus was thicker for SMA than for M1, indicating that SMA encodes relatively more information about future position (Fig. 4E-F). Consistent with this, mutual information about future position, relative to that about current position, fell off more steeply with prediction horizon in M1 than in SMA (Fig. 4G), in each monkey (*p <* 0.001; repeated-measures ANOVA). Since SMA is not more sensitive than M1 to small changes in future position, this difference must come from how well SMA distinguishes future positions that are far apart.

### The multi-scale geometry reflects the statistics of natural stimuli

The Fisher information depends only on how neurons respond to stimuli. The mutual information, and by extension the multi-scale geometry, depends both on how neurons respond to stimuli and on the stimulus distribution itself. Intuitively, a population that is sensitive only to stimulus changes that rarely occur will convey little information.

To illustrate this, we considered a two-dimensional stimulus whose distribution is concentrated along a narrow, curved ridge (Fig. 5A). This mimics natural images, which are thought to lie close to a low-dimensional manifold within the space of all possible images^49–52^. For a neural code that encodes both stimulus dimensions with equal sensitivity, the Fisher geometry is uniform, with no trace of the stimulus distribution (Fig. 5B). The multi-scale geometry, in contrast, is stretched along the ridge, where stimulus variations are common (Fig. 5C).

**Figure 5.**
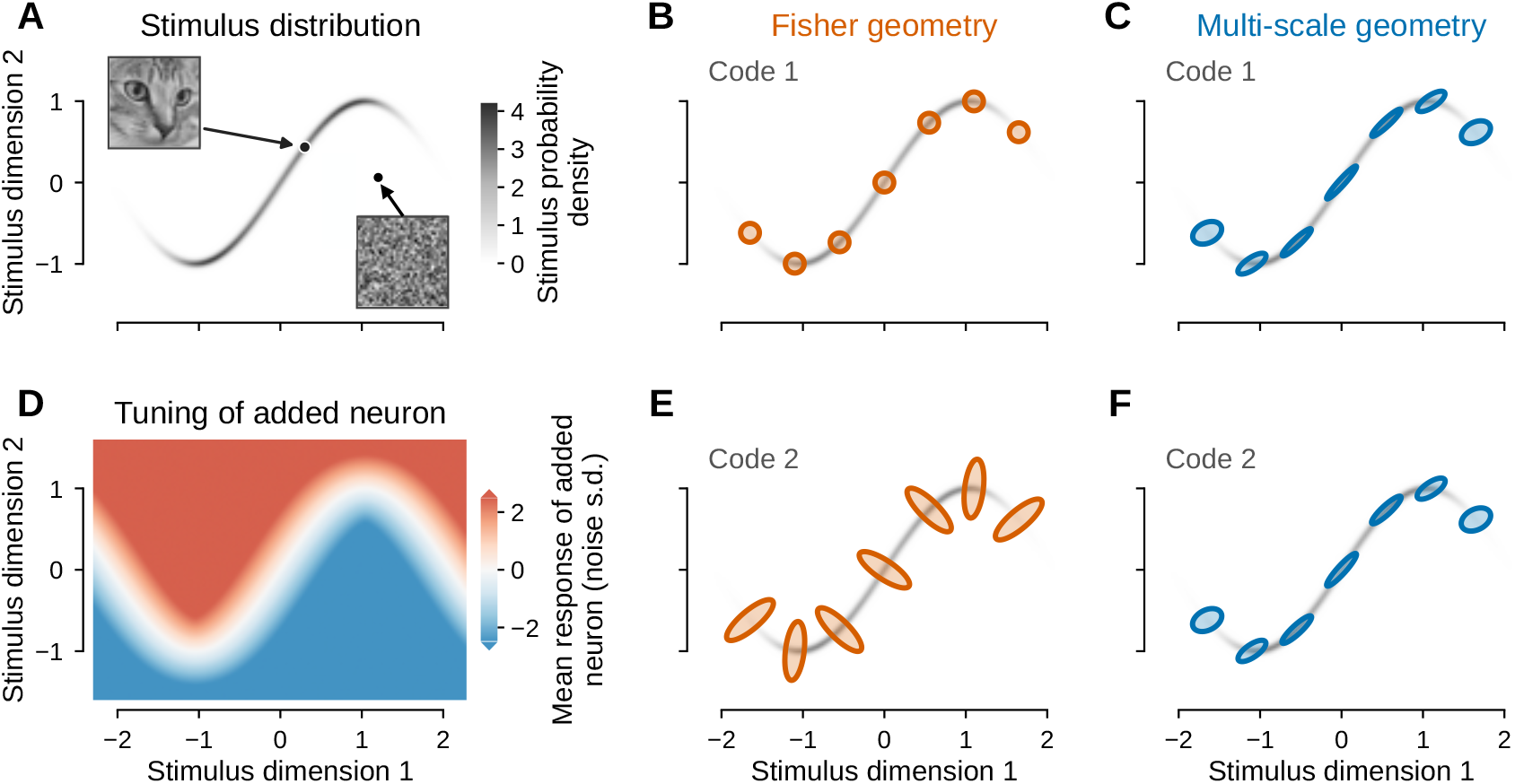
The multi-scale geometry reflects the stimulus distribution. (A)Distribution of a two-dimensional stimulus, concentrated along a narrow ridge. Example images are illustrative: stimuli on the ridge correspond to natural images, which have high probability of occurring, and stimuli away from it to images that occur with vanishingly small probability. (B)Fisher geometry for a code that encodes both stimulus dimensions with equal sensitivity (code 1). Ellipses show the metric tensor at points along the ridge, corresponding to how much the geometry stretches distances along each direction. (C)Multi-scale geometry for the same code, which is stretched along the ridge of the prior. (D)Tuning of a neuron added to form code 2. It responds in proportion to the distance of the stimulus from the ridge, and is almost silent for stimuli drawn from the distribution. (E)Fisher geometry for code 2, dominated by sensitivity to stimulus changes away from the ridge. (F)Multi-scale geometry for code 2, almost unchanged from (C).

We then added a single neuron that responds only when the stimulus leaves the ridge (Fig. 5D). This neuron is almost silent for stimuli that actually occur. Yet it transforms the Fisher geometry, which becomes dominated by sensitivity to changes orthogonal to the ridge, which rarely occur (Fig. 5E). The multi-scale geometry, by contrast, is barely changed, since the added neuron conveys almost no information (Fig. 5F). Thus, two codes that convey nearly the same information can have very different Fisher geometries, if one of them is sensitive to stimulus changes that rarely occur. As we show below, this has practical consequences when analysing neural responses to natural images.

### The multi-scale geometry reveals which image features are encoded in V1 and V4

We next asked what the multi-scale geometry reveals about how neurons in visual cortex encode natural images. The rich, high-dimensional structure of natural images makes this a hard problem, and has long limited information-theoretic analyses of neural responses to natural stimuli^53^. To estimate the multi-scale geometry, we need the neural sensitivity at each scale, ***J*** _*t*_, which requires an integral over stimuli (Eq. 3) that is intractable for high-dimensional, non-Gaussian stimuli and responses. We overcame this using diffusion models^54–56^, the generative models behind modern image synthesis, which we trained to sample from the posterior distribution of images given neural responses, *p*(***x*** | ***r***). Remarkably, in learning to do so, a diffusion model implicitly learns how this posterior density varies across image space, in a form from which the multi-scale geometry can be computed directly^37,43^ (Supplementary note).

We analysed the dataset of Papale et al.^57^, consisting of large-scale electrophysiological recordings from V1 and V4 of awake macaques viewing natural images, and selected the 50 most reliable neurons from each area. Estimating the geometry requires knowing how neurons would respond to images that were never shown in the experiment. We therefore first fitted a model of how these neurons respond to images, *p*(***r*** | ***x***) (Fig. 6A). We used a deep convolutional neural network (Inception architecture^58^; Methods), which achieved 67% and 63% of explainable variance in V1 and V4, respectively, comparable to the encoding models reported for these data by Papale et al.^57^ (Fig. 6B–C). We then trained a diffusion model using the responses of this network, and used it to estimate the multi-scale geometry.

**Figure 6:**
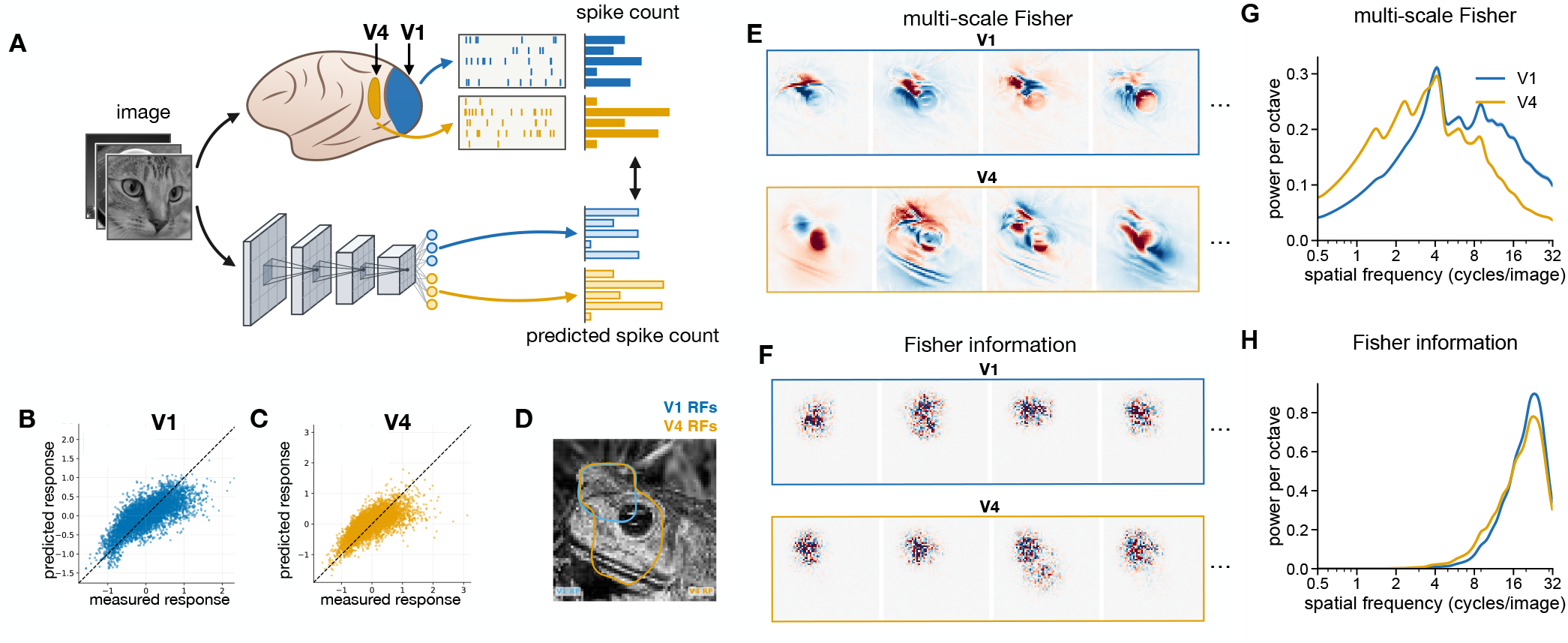
Multi-scale geometry of neural population recordings in V1 and V4. (A)We trained a deep convolutional network to predict the spiking responses of V1 and V4 neurons to natural images. (B)Measured versus predicted responses across all V1 neurons and images. (C)Same as in (B), for V4. (D)Example image, with the regions covered by V1 and V4 receptive fields outlined in blue and yellow, respectively. (E)First four eigenvectors of the multi-scale metric tensor for this image, shown for V1 (top) and V4 (bottom). These identify stimulus variations that contribute most to encoded information. In V1, features are localised and edge-like. In V4, features are broader and more global, with the leading eigenvector capturing the overall shape of the stimulus. (F)Same as in (E), but for the Fisher geometry. Features in V1 and V4 are qualitatively similar and dominated by high-frequency noise, making them hard to interpret in terms of encoded stimulus features. (G)Power spectrum of the leading ten eigenvectors of the multi-scale metric tensor, averaged over 200 held-out test images (shading, ±s.e.m. across images). Power is shown per octave, so the area under each curve gives the fraction of power in that frequency range. V1 eigenvectors carry more of their power at high spatial frequencies than V4 eigenvectors (median half-power frequency 5.0 vs 2.8 cycles/image; higher in V1 in 199 of 200 images, sign test *p <* 0.001). (H)Same as in (G), for the Fisher geometry. In both areas almost all power lies at high frequencies, near the pixel resolution, and the two spectra differ by about a tenth of an octave (median half-power frequency 20.0 vs 18.6 cycles/image; higher in V1 in 196 of 200 images, *p <* 0.001).

The multi-scale geometry identifies, for each reference image, the changes to that image that the population encodes best. These are the leading eigenvectors of ***G***(***x***): the directions along which image space is most strongly stretched, analogous to the major axes of the ellipses in Fig. 1A. Each eigenvector can be displayed as a pattern of pixel changes, showing a change to the reference image that contributes strongly to the encoded information. For a reference image of a frog (Fig. 6D), the leading eigenvectors differed between V1 and V4 (Fig. 6E; see Fig. S1 for further reference images). In V1, they emphasised edges and fine detail. In V4, they followed more global structure, such as the shape of the frog’s eye. To quantify this difference, we computed the spatial frequency content of the leading eigenvectors for each reference image. V1 eigenvectors had relatively more power at high spatial frequencies than V4 eigenvectors for almost every reference image (Fig. 6G). This is consistent with the known hierarchy of visual processing, in which V4 neurons are selective for more global shape features than V1 neurons^59^. The Fisher geometry gave a very different picture. Its leading eigenvectors resembled high-frequency noise, confined to the neurons’ receptive fields (Fig. 6F). In both areas, almost all of the eigenvectors’ power lay at high spatial frequencies, and their power spectra differed only slightly between V1 and V4 (Fig. 6G–H). The Fisher geometry thus revealed no interpretable image features, and only a weak difference between the two areas. We attribute this to the effect illustrated in Fig. 5. Adding a noise-like pattern to a natural image produces an image that would essentially never occur in nature, like a stimulus off the ridge in Fig. 5A. Because an encoding model is fitted only to responses to natural images, it can be highly sensitive to such changes without any cost to how well it fits the data. The Fisher geometry is then dominated by this sensitivity (as in Fig. 5E). The multi-scale geometry is not, since changes that almost never occur contribute almost nothing to the encoded information (as in Fig. 5F).

This explanation makes a testable prediction. A model’s sensitivity to image changes that essentially never occur is not constrained by the data, so two models that fit the same data equally well have no reason to share it. Their Fisher geometries should therefore differ, while their multi-scale geometries should agree. To test this, we fitted a second encoding model, with a different architecture (ResNet^60^, rather than the inception model used above; Methods), to the same recordings (Fig. 7A). The two models predicted V1 and V4 responses equally well (Fig. 7B), and their multi-scale geometries were similar (Fig. 7C). Across reference images, the multi-scale geometries of the two models were more closely aligned when fitted to the same area than when fitted to different areas (Fig. 7E). For the Fisher geometry, the reverse was true. It differed markedly between the two models (Fig. 7D), and was more closely aligned between different areas fitted with the same model than between the two models fitted to the same area (Fig. 7F). The differences between V1 and V4 revealed by the multi-scale geometry therefore do not depend on the choice of model. As predicted, the Fisher geometry reflected the choice of model more than the neural data, whereas the multi-scale geometry reflected the data.

**Figure 7:**
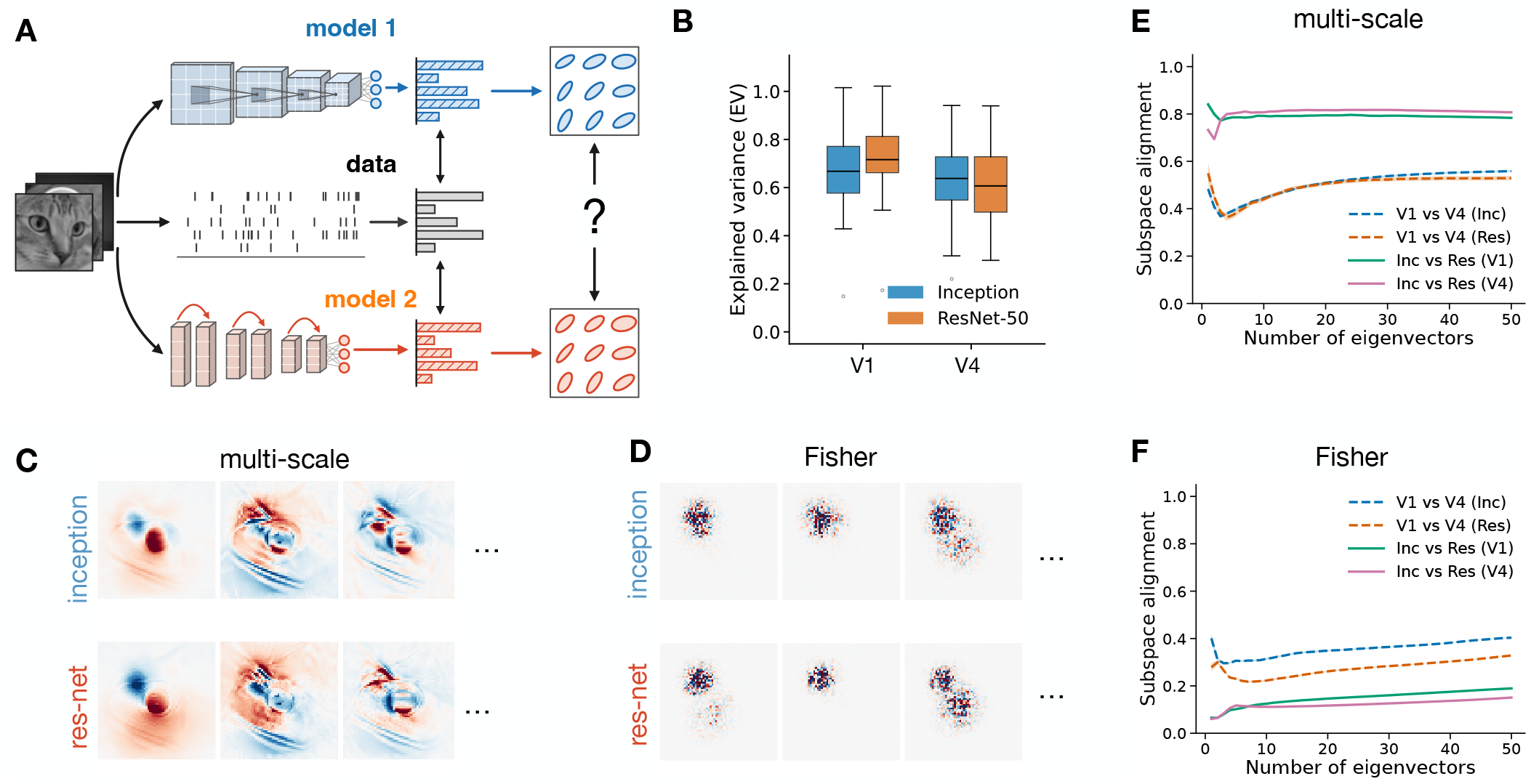
Multi-scale geometry is robust to model choices. (A)We fitted two encoding models with different architectures (Inception and ResNet-50; see Methods) to the same recordings, and compared the geometries they induce. Since both models are fitted to the same data, any difference between their geometries reflects the choice of model architecture. (B)Fraction of explainable variance predicted by each model on held-out test images, for the 50 V1 and 50 V4 neurons (boxes, quartiles; lines, medians; whiskers, 1.5 times the interquartile range). The two models predict the responses about equally well: ResNet-50 is slightly better in V1 (mean 0.73 vs 0.67; Wilcoxon signed-rank test across neurons, *p <* 0.001) and Inception slightly better in V4 (0.63 vs 0.62; *p* = 0.04). (C)First three eigenvectors of the multi-scale metric tensor for the example image in Fig. 6D, computed from the Inception (top) and ResNet-50 (bottom) models fitted to V4. The two models yield qualitatively matching eigenvectors, identifying the same stimulus variations as contributing most to encoded information. (D)Same as in (C), for the Fisher geometry. The detailed spatial structure of the eigenvectors differs markedly between the two models. (E)Subspace alignment of the leading *k* eigenvectors of the multi-scale metric tensor, against *k*, averaged over 200 held-out test images. Alignment between different models fitted to the same area (solid) is consistently higher than between the same model fitted to different areas (dashed); for *k* = 10 this holds in all 200 images (Wilcoxon signed-rank test across images, *p <* 0.001). (F)Same as in (E), for the Fisher geometry. The trend is reversed in all 200 images (*p <* 0.001): alignment is higher between the same model fitted to different areas than between different models fitted to the same area, indicating that the Fisher geometry depends more on the choice of model than on the neural data.

## DISCUSSION

Over half a century ago, Barlow’s efficient coding hypothesis proposed that sensory systems are adapted to transmit as much information as possible about the environment, subject to constraints such as noise and metabolic cost^38,40,61^. More recently, representational geometry has become a central tool for describing what neural populations encode^5,6,62,63^. Yet these two lines of work have remained largely separate, since no representational geometry has been tied directly to mutual information. The multi-scale geometry provides this link: its distances are exactly tied to mutual information, so an informative code is one that expands stimulus space. Moreover, while mutual information gives a single number, the geometry shows how different stimulus features contribute to the total. For example, it reveals how neurons with heterogeneous tuning combine to encode information (Figs. 2–3), how different motor areas encode upcoming movements (Fig. 4), the effect of the stimulus distribution (Fig. 5), and which visual features are encoded by neurons in V1 and V4 (Figs. 6–7).

The geometry of a neural code also constrains perception. The Fisher information of a sensory population bounds fine discrimination, predicting perceptual thresholds from the properties of the neurons themselves (Fig. 1)^42,64^. In colour vision, for example, the Fisher information of the cone photoreceptors predicts how finely colours can be discriminated^65^, and thus the perceptual geometry of colour space^66–68^. Likewise, since information can only be lost through processing, the multi-scale distances between stimuli measured from neural responses place an upper bound on those measured from an animal’s perceptual judgements. Where perceptual distances approach this bound, the animal exploits essentially all the information in the recorded population; where they are shorter, the comparison reveals which stimulus features the neurons encode but the animal fails to use.

Previous work showed that, in the high signal-to-noise limit, the mutual information can be approximated using the average log-determinant of the Fisher information^69^. This relation has motivated work deriving how efficient codes, which maximise information transfer, depend on the stimulus distribution^41,70^. Combined with Bayesian decoding, such codes predict systematic biases in human perception and their lawful relation to discrimination thresholds^64,71,72^. However, this approximation can fail catastrophically, and even diverge, at low signal-to-noise or when there are few neurons per stimulus dimension^35,36,73^. Yet these regimes matter: recorded populations are often small, while the very perceptual biases that efficient coding explains would vanish if coding were near-perfect^71^. In contrast, the relation between the multi-scale geometry and mutual information is exact at any signal-to-noise, with the above approximation of Brunel and Nadal^69^ recovered from it as a limiting case. This opens the way to deriving optimal codes, and the perceptual biases they predict, beyond the high signal-to-noise limit.

A basic question about any neural population is which features of a stimulus it encodes. Recent work answers this for single neurons by synthesising the image that drives a neuron most strongly, its most exciting input^74–76^. Meanwhile, an earlier, information-theoretic approach looks for the few linear projections of the stimulus that carry the most information about a neuron’s spikes, its maximally informative dimensions^53^. However, both these approaches make strong assumptions about neural encoding: in the former, that a neuron’s preferred stimulus summarises what it encodes; in the latter, that neurons encode a few linear projections of the stimulus. Moreover, both approaches characterise neurons one at a time. Extending them to a population means choosing how to combine neurons’ responses into a single objective, such as driving one neuron while suppressing the others^75^, and it is not clear which choice reflects what the population encodes as a whole.

Another way to ask which features a population encodes is to find the changes to a given stimulus that most alter its responses, as measured by the gradient of neural tuning curves^77^ or by their Fisher information^16,33,78,79^. For high-dimensional stimuli such as images, the leading eigenvectors of the Fisher information give the stimulus changes to which the population is most sensitive. However, our results show that this measure can be dominated by the encoding model’s sensitivity to image changes that essentially never occur in natural images. Because such changes are absent from the data, a fitted model can be highly sensitive to them without fitting the data any worse. In V1 and V4, the leading eigenvectors of the Fisher information thus resembled high-frequency noise, and varied greatly between different models fitted to the same data. The multi-scale geometry discounted these image changes, picking out only those that contribute to encoded information (Fig. 5). The resulting features were interpretable, and differed between the two areas: edges and fine detail in V1, more global structure in V4 (Fig. 6). They were also stable across different models fitted to the same data (Fig. 7).

A longstanding objection to information-theoretic approaches in neuroscience is that mutual information is prohibitively hard to estimate for high-dimensional natural stimuli and large populations^80,81^. This has often confined information-theoretic analyses to low-dimensional or artificial stimuli^30,53^. Diffusion models offer a way around this^43,82,83^. A model trained to sample images from the posterior, *p*(***x*** | ***r***), implicitly learns how that density varies across image space. This is exactly what the multi-scale geometry requires. We could therefore measure how much information populations in V1 and V4 encode about natural images, and how different image features contribute, despite the complex, high-dimensional structure of these stimuli (Fig. 6-7).

Comparing population codes through the geometry of their responses is now routine, across areas, tasks, species and models^7,12,20,22,62^. What such comparisons tell us, however, depends on how the geometry is defined. Most previous geometries rest on choices about which aspects of the response to take into account, such as ignoring response variability^20–22^, using only mean responses and covariances^14,26,28^, or measuring distances in the latent space of a deep network^84,85^. Like the Fisher information, the multi-scale geometry makes no such choices about which response statistics to include, or how the neural code is read out. But where the Fisher geometry reflects only local sensitivity, which governs fine discrimination, the multi-scale geometry reflects how well neurons distinguish stimulus changes at every scale. Multi-scale distances are tied exactly to mutual information, and can only shrink under further processing, bounding what any downstream circuit could extract about the stimulus. A comparison between two areas, or between a brain and a model, thus becomes a comparison of the information each code carries. Computing the geometry requires only a model of how neurons respond to stimuli, together with samples of the stimuli themselves. As such models become available for more brain areas, species and sensory modalities^86–89^, the multi-scale geometry offers a way to map what each encodes about the natural world.

## METHODS

### Estimating the multi-scale geometry for low-dimensional stimuli

For low-dimensional stimuli (Figs 1-5) we estimated ***G***(***x***) by combining Monte Carlo sampling with numerical integration over a grid of stimulus values. Combining Eqs. 2 and 4, the multi-scale Fisher information matrix is

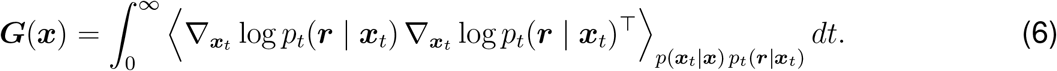

To estimate this integral, we changed the integration variable to log *t* and bounded it to a finite range of scales:

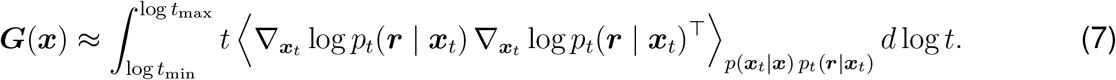

We evaluated this by Monte Carlo sampling. Each sample was generated in three steps: we drew a scale *t* uniformly in log *t* over [*t*_min_, *t*_max_]; a blurred stimulus ***x***_*t*_ ~ *p*(***x***_*t*_ | ***x***) = *N*(***x***, *t****I***); and a response ***r*** ~*p*_*t*_(***r*** | ***x***_*t*_). The integrand of Eq. 7 was then averaged over *N* samples and multiplied by log(*t*_max_*/t*_min_).

Responses were drawn by ancestral sampling, following Eq. 3: we first drew a stimulus ***x***′~*p*(***x*** |***x***_*t*_), then a response ***r*** ~*p*(***r*** |***x***′). The gradient in the integrand follows from differentiating Eq. 3 with respect to ***x***_*t*_:

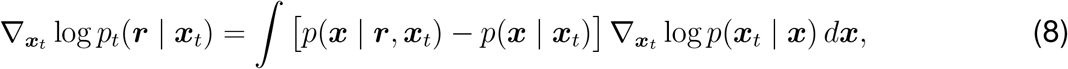

where *p*(***x*** | ***x***_*t*_) ∝*p*(***x***) *p*(***x***_*t*_ | ***x***) and *p*(***x*** | ***r, x***_*t*_) ∝*p*(***r*** | ***x***) *p*(***x*** | ***x***_*t*_). For Gaussian blur, 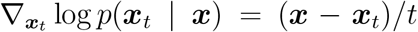. All integrals and normalisations over stimuli were computed numerically on a grid of stimulus values, and ***x***′ was drawn from the resulting discrete distribution. The procedure thus requires only the stimulus prior and an encoding model whose likelihood, *p*(***r*** | ***x***), can be evaluated and sampled.

For the model populations in Figs. 1, 2 and 5, we assumed Gaussian noise with covariance **σ**, so that log 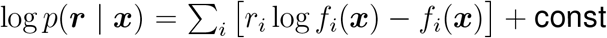 const, where ***f*** (***x***) are the tuning curves. For the grid-cell recordings in Fig. 3, we assumed independent Poisson noise, so that log *p*(***r*** |***x***) = σ_*i*_ [*r*_*i*_ log *f*_*i*_(***x***) − *f*_*i*_(***x***)] + const, where *r*_*i*_ is the spike count of neuron *i* and *f*_*i*_(***x***) its tuning curve, expressed as the expected count in a time bin and estimated from the data. For the motor-cortex recordings in Fig. 4, the response at each moment of a movement was the trial-averaged population state, summarised by its first eight principal components, plus isotropic Gaussian noise with variance *σ*^2^; since the same stimulus occurred at several moments, its likelihood was the mixture of the Gaussians centred on the states at those moments.

For a periodic stimulus dimension, such as grating orientation in Fig. 2 and each of the two pedal angles in Fig. 4, the only change is that blurring wraps around the circle. We used a wrapped Gaussian, *p*(*x*_*t*_| *x*) ∝ σ_*ℓ*_ exp (−(*x* −*x*_*t*_ + 2*πℓ*)^2^*/*2*t*, where the sum runs over the integer number of turns *ℓ* around the circle, truncated at |*ℓ*|≤ 5, with *x*_*t*_ wrapped onto [−*π, π*). The gradient required in Eq. 8 is then

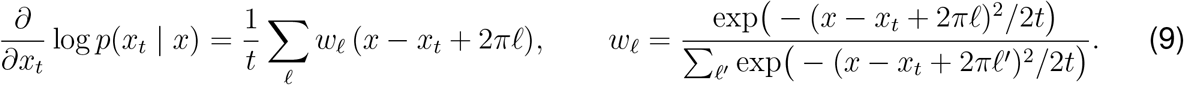

For the pedal angles of Fig. 4, measured in cycles, 2*πℓ* is replaced by *ℓ*, and the two angles were blurred independently with the same *t*, so that blurred stimuli stay on the torus. Because the scales there reach *t* = 4 cycles^2^, we evaluated the wrapped sum with |*ℓ*| ≤ 3 for *t <* 0.15 and, for larger *t*, through the Fourier series of the wrapped Gaussian in the angle difference,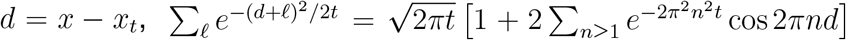 (an exact identity), truncated at *n* = 6; the factor 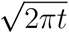 cancels in the posterior.

### Estimating the multi-scale geometry for high-dimensional stimuli

For natural images, the integral over stimuli in Eq. 3 is intractable, so the gradient in Eq. 8 cannot be computed on a grid. We therefore used the diffusion-model estimator of Azeglio et al.^37^, which builds on Laquitaine et al.^43^; we summarise it here and refer to that work for derivations. From Eq. 6, squared distances along any image direction ***dx*** are

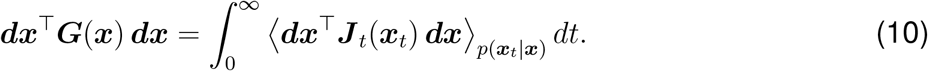

In Azeglio et al. ^37^ we show that applying Bayes’ rule and Tweedie’s formula^90^ to the gradient of log *p*_*t*_(***r*** | ***x***_*t*_), the sensitivity at each scale can be written as

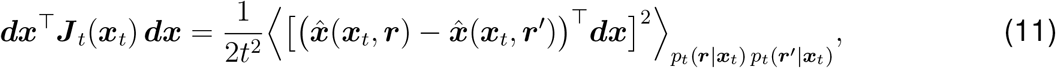

where 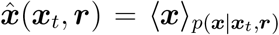 is the best estimate of the image given the blurred image and the neural response, and ***r*** and ***r***′ are two independent responses. The estimate 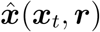 is exactly what a conditional diffusion model learns when trained to denoise images given neural responses^55^. Details of the encoding model *p*(***r***| ***x***) and denoising diffusion model, for estimating 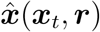 are given later, when we describe the simulations performed to produce Figs 6-7.

### One-dimensional tuning curves (Fig. 1)

We considered a single neuron encoding a scalar stimulus *x*, drawn from a uniform prior on [−7, 7], with Gaussian response noise, *p*(*r* | *x*) = *N* (*f* (*x*), *σ*^2^). The three tuning curves were

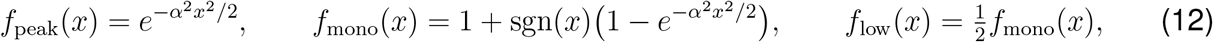

with *α* = 1.8. By construction, 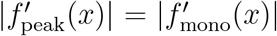 for all *x*. The noise variance was *σ*^2^ = 0.03 for the single-peaked and monotonic neurons, and *σ*^2^ = 0.05 for the low-gain neuron. Stimulus space was discretised on a grid of 5,000 points.

The multi-scale metric *G*(*x*) was estimated as described above, at 500 equally spaced reference stimuli between *x* = −3 and *x* = 3, using *N* = 30,000 Monte Carlo samples and scales *t* ∈ [10^−4^, 300]. The Fisher information was *J* (*x*) = *f* ′(*x*)^2^*/σ*^2^. Distances from the reference stimulus *x*_0_ = −3 were computed by integrating the line element,

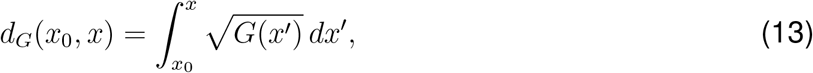

and likewise for the Fisher geometry with *J* in place of *G*. Fine discrimination performance, shown as *d*′ per unit stimulus difference, was 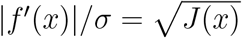.

Mutual information was estimated as

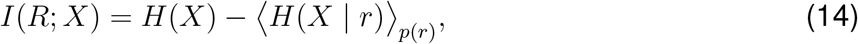

where *H*(*X*) is the entropy of the uniform prior and *H*(*X* | *r*) the entropy of the posterior. We drew 30,000 stimulus–response pairs, *x*~ *p*(*x*) and *r* ~ *p*(*r*| *x*), computed each posterior on the stimulus grid by Bayes’ rule, *p*(*x*| *r*) ∝ *p*(*r* | *x*) *p*(*x*), and evaluated its entropy by numerical integration over the grid.

### Simple model of orientation and contrast encoding in primary visual cortex (Fig. 2)

Stimuli were gratings described by their orientation, *θ*, mapped onto [− *π, π*), and their signed contrast, *c*, whose sign gives the grating polarity. Both were drawn from a uniform prior, with contrast scaled to [−*π, π*] so that blurring treats the two dimensions equally; figures show contrast normalised to [−1, 1]. The complex-cell population consisted of 10 neurons with preferred orientations *θ*_*i*_ evenly spaced on [−*π, π*) and tuning curves

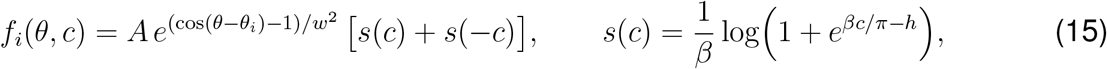

where *s* is a smooth rectification. Parameters were *A* = 0.6, *w* = *π/*4, *β* = 50 and *h* = 0.1. The simple-cell population was obtained by splitting each complex cell into a pair, with tuning curves

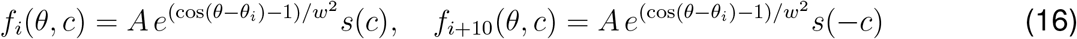

Stimulus space was discretised on a 200 × 200 grid.

The multi-scale metric was estimated at 2,000 reference stimuli drawn from the prior, using *N* = 10,000 Monte Carlo samples and scales *t* ∈ [0.005, 100], with orientation treated as a periodic dimension. To visualise each geometry, we computed geodesic distances between reference stimuli as shortest paths on a 50-nearest-neighbour graph, with each edge weighted by its length under the metric, computed with the average of the metrics at its two ends, and embedded them in three dimensions by multidimensional scaling with a relative stress, σ_*i<j*_ (∥***y***_*i*_− ***y***_*j*_ ∥*/d*_*ij*_− 1)^2^, so that short and long distances are weighted equally. The embedding was initialised by standard metric multidimensional scaling (SMACOF^91^) and then refined by gradient descent (Adam^92^) on the relative stress. Here *d*_*ij*_ is the geodesic distance between reference stimuli *i* and *j*, and ***y***_*i*_ the embedded position of stimulus *i*. Mutual information was estimated as for Fig. 1.

### Population coding by grid cells in entorhinal cortex (Fig 3)

We used the recordings of Gardner et al.^48^ from rat R, day 1, comprising 483 grid cells in three modules (166, 168 and 149 cells), restricted to the open-field session. The animal’s position at each spike time was obtained by interpolating the tracked position. Spike counts and occupancy time were binned on a 50 × 50 grid spanning the arena, each smoothed with a Gaussian kernel of width 1.5 bins, and divided to give a firing-rate map; bins with less than 10 ms of smoothed occupancy were set to zero. Positions were rescaled to the unit square, and each cell’s tuning curve, *f*_*i*_(***x***), was obtained by bilinear interpolation of its rate map, with a floor of 0.01 Hz, and expressed as the expected spike count in a 100 ms time bin.

We chose two populations of 20 cells to illustrate how the two geometries can differ. The fixed-spacing population was chosen to be highly sensitive to small changes in position: we ranked cells by their Fisher information averaged over the arena, 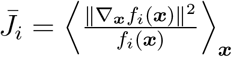, and took the 20 highest-ranked cells from the module with the finest spacing. The mixed-spacing population was chosen to carry more information: starting from the fixed-spacing population, at each of 180 steps a randomly chosen cell was replaced by a randomly chosen candidate from the 20 highest-ranked cells of each of the other two modules, and the swap was kept if it increased the mutual information. The resulting population contained 7, 8 and 5 cells from the three modules. Once the populations were selected, the Fisher and multi-scale geometries were computed in exactly the same way for both, so the comparison between the two geometries is unaffected by how the cells were chosen.

The multi-scale and Fisher metrics were computed for each population as described above, with a uniform prior over the arena discretised on a 400 × 400 grid, 1,000 reference positions drawn from the prior, *N* = 5,000 Monte Carlo samples and scales *t* ∈ [10^−5^, 100]. Each geometry was embedded in two dimensions by multidimensional scaling, as for Fig. 2, using a 20-nearest-neighbour graph, and rotated to align with the arena. Mutual information in Fig. 3E was estimated by Monte Carlo, as for Fig. 1.

### Coding of movement trajectories in motor areas (Fig 4)

We used the recordings of Russo et al.^8^ from two monkeys (C and D) turning a hand pedal forward or backward through 0.5 to 7 cycles (20 movement conditions). Trial-averaged firing rates at 1 ms resolution were soft-normalised and centred, and projected onto their first eight principal components, separately for each area and monkey as performed by Russo et al.^8^. Each movement was analysed from 100 ms before its onset to 100 ms after its end.

The stimulus at time *τ* was the pair of pedal angles (*θ*_*τ*_, *θ*_*τ*+*δ*_), in cycles, for prediction horizons *δ* from 100 to 1,000 ms, sampled every 10 ms while *τ* + *δ* stayed within the same movement. Both angles were divided into 48 bins per cycle, so that moments falling in the same pair of bins were the same stimulus, and the prior was uniform over moments. Responses were the population state with isotropic Gaussian noise, of standard deviation 10% of its root-mean-square amplitude.

The multi-scale metric was computed from Eq. 7, with a wrapped Gaussian blur on each angle, the posterior computed over the sampled moments, *N* = 24,000 Monte Carlo samples and *t* ∈ [10^−6^, 4], with scales below *t*_min_ approximated from ***J***. It is a 2 × 2 matrix, ***g***, in the coordinates of the two angles, and 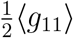 and 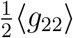 are the information carried along the current and the future angle (Eq. 5). The Fisher information, ***J*** = *σ*^−2^(*∂****f*** */∂****x***)^⊤^(*∂****f*** */∂****x***), was computed with the chain rule, 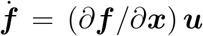, where 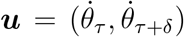 and both rates were estimated from 1 ms differences. The population state is known only along the path that each movement traces on the torus, so this gives ***J*** only along that path, and ***J*** is therefore of rank one, 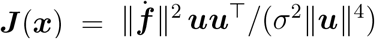; it was set to zero while the pedal was still. Averages over stimuli used 800 moments evenly spaced across all movements.

For Fig. 4B–C and E–F, the metrics of the two monkeys were averaged in each of the 48 × 48 cells of the torus (for ***J***, over the monkeys that visited the cell). The cells were then embedded in three dimensions by multidimensional scaling with the same relative stress as for Fig. 2: geodesic distances were computed as shortest paths on the periodic grid, each cell linked to its eight neighbours, and the stress was minimised by SMACOF. The Fisher metric is of rank one at visited cells and zero elsewhere, so many of its distances would otherwise be zero. For the embedding only, we therefore added 0.1 times the mean of 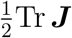 over visited cells to its diagonal; the circles and ratios used the Fisher metric without it. The circles have circumferences proportional to the square roots of the two diagonal elements of each metric, averaged over moments and monkeys, on the same scale for both areas. Because they split the joint information between two correlated angles, their ratio can differ slightly from that in Fig. 4G.

In Fig. 4D and G, the Fisher ratio was ⟨ *J*_22_ ⟩ */* ⟨ *J*_11_⟩ and the information ratio *I*(*R*; *θ*_*τ*+*δ*_)*/I*(*R*; *θ*_*τ*_), with each mutual information estimated by Monte Carlo as for Fig. 1, with one binned angle as the stimulus. The panels show the mean of the two monkeys’ ratios, with standard errors obtained per monkey by resampling movements (bootstrap for the Fisher ratio, leave-one-movement-out jackknife for the information ratio) and combined across monkeys as independent. The areas were compared for each monkey and ratio with a repeated-measures ANOVA with factors area and horizon, using as subjects the movements present at every horizon (15 in monkey C, 16 in monkey D), each contributing its jackknife pseudo-value. The interaction term was Greenhouse– Geisser corrected.

### Multi-scale geometry and stimulus prior (Fig 5)

Stimuli were two-dimensional, ***x*** = (*x*_1_, *x*_2_), and distributed along a narrow curved ridge. We wrote each stimulus as

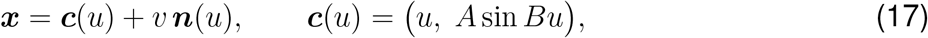

where ***c***(*u*) is the centre line of the ridge, ***n***(*u*) its unit normal, *u* the position along the ridge and *v* the signed distance from it. Position along the ridge was drawn from *q*(*u*) ∝ exp (−*u/L*)^4^, and distance from it from *N* (0, *s*^2^), with *A* = 1, *B* = 1.5, *L* = 1.35 and ridge width *s* = 0.03.

In the first code, two neurons encoded the two stimulus dimensions with equal sensitivity, *r*_*j*_ = *x*_*j*_ + *σξ*_*j*_ for *j* = 1, 2, with *ξ*_*j*_~ *N* (0, 1) and *σ* = 0.5. The second code added a third neuron whose mean response was proportional to the signed distance of the stimulus from the ridge, *r*_3_ = *k d*(***x***) + *σξ*_3_, with *k* = 3. Since *d*(***x***) is zero along the ridge, this neuron’s mean response to stimuli drawn from the distribution is about *ks* = 0.09, roughly 0.2 noise standard deviations; Fig. 5D shows *k d*(***x***)*/σ*, with the colour scale saturating at±2.5. The Fisher information is ***J*** = *σ*^−2^***I*** for the first code and ***J*** (***x***) = *σ*^−2^ (***I*** + *k*^2^ ∇*d* ∇*d*^⊤^) for the second. The multi-scale metric was estimated as described above, at points equally spaced in *u* along the centre line.

Ellipses show the metric tensor at a subset of these points. Each ellipse is centred on the point, with principal axes along the eigenvectors of the metric and semi-axes proportional to the square roots of its eigenvalues, so that it is elongated along the stimulus directions that the geometry stretches most. Within each geometry, the same scale was used for both codes, so that ellipses can be compared directly between the top and bottom rows.

### Multi-scale geometry of neural population recordings in V1 and V4 (Figs. 6–7)

#### Encoding models

We analysed the recordings of Papale et al.^57^, from V1 and V4 of a macaque viewing 22,248 natural images from the THINGS database^93^. For each area we selected the 50 channels with the highest reliability across odd and even trials, and fitted their trial-averaged normalised multi-unit responses, ***f*** (***x***), to grayscale 64 × 64 versions of the images. Each encoding model consisted of the early layers of a network pretrained on ImageNet, kept fixed, followed by a spatial readout for each channel, a Gaussian-weighted sum of the network’s features, fitted by minimising the mean-squared error on the training images. The receptive-field regions outlined in Fig. 6D are the union over the 50 channels of the region where each channel’s Gaussian readout weight exceeds half its maximum, mapped from the feature map onto the image; they do not include the receptive fields of the fixed network layers. We used InceptionV3^58^ up to layer Conv2d 4a 3×3 and, for Fig. 7, ResNet-50^60^ up to layer layer1.0. Predictive performance was measured on the held-out test images of Papale et al. The mean correlation between predicted and recorded responses of the selected neurons was 0.75 (inception) and 0.78 (ResNet) in V1, and 0.71 and 0.70 in V4; Papale et al.^57^ report 0.75–0.76 in V1 and 0.74– 0.76 in V4, averaged over all reliable recording sites of both monkeys. Responses were modelled as *p*(*r*|*x*) = *N* (***f*** (***x***), *σ*^2^***I***), with *σ* = 0.22 matched to the median trial-to-trial variability of the recorded channels.

#### Diffusion model

For each encoding model, we trained a conditional diffusion model^55^ to estimate 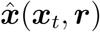 in Eq. 11. Training used 100,000 grayscale 64 × 64 images from Tiny ImageNet^94^, each paired with a response sampled from the encoding model, ***r*** = ***f*** (***x***) + *σ****ξ*** with ***ξ*** ~ *N* (**0, *I***), redrawn at every training step. The network was a U-Net, a standard convolutional architecture for image denoising, which also received the response as input. On a random 10% of training steps the response was withheld, so that the same network also learned the estimate given the blurred image alone, 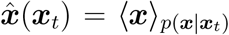^95^. Architecture and training details are given in Azeglio et al.^37^.

We evaluated Eq. 11 at 24 scales, *t*_*j*_, ranging from *t* = 0.02 to 1.1 × 10^4^ for images scaled to [− 1, 1]. At each scale we drew *N* = 200 blurred images ***x***_*t*_ ~ *N* (***x***, *t*_*j*_***I***) and, for each, two responses ***r, r***′. Exact sampling from *p*_*t*_(***r*** |***x***_*t*_) would require drawing images from *p*(***x***| ***x***_*t*_) by running the diffusion model’s full generative process for every sample. Following previous work^43^, we instead drew both responses from 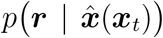, an approximation that is most accurate at fine scales, where *p*(***x*** | ***x***_*t*_) is concentrated around its mean.

Since images contain *P* = 64^2^ pixels, we never formed the *P* ×*P* matrix ***G***(***x***) explicitly. Instead, each Monte Carlo sample contributed one row to a matrix ***D***,

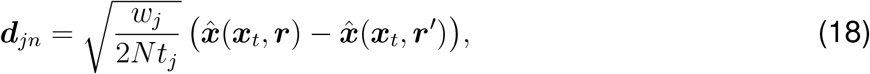

so that summing over scales approximates the integral in Eq. 7, and ***G***(***x***) ≈ ***D***^⊤^***D***. The leading eigenvectors and eigenvalues of ***G***(***x***) then follow from the singular value decomposition of ***D***, which is much smaller than ***G***(***x***); we kept the leading 50. Similarly, the eigenvectors of ***J*** (***x***) = *σ* ^−2^ ∇_***x***_***f*** ^⊤^ ∇_***x***_***f*** follow from the singular value decomposition of the Jacobian ∇_***x***_***f*** (***x***), so ***J*** (***x***) has at most 50 non-zero eigenvalues, one per channel. All analyses used 200 held-out images.

#### Spatial frequency content

For Fig. 6G–H, each of the leading ten eigenvectors was normalised to unit power, and its two-dimensional power spectrum was summed over rings of one cycle per image to give power as a function of spatial frequency; the zero-frequency component, a uniform change of the whole image, falls outside the rings. For each image, we averaged the spectra of its ten eigenvectors. For display, the power in each ring was spread uniformly over the ring’s range of log2 frequency and smoothed with a Gaussian of standard deviation 0.08 octaves, so that the area under the curve on a logarithmic frequency axis gives the fraction of power in each frequency range. The half-power frequency of an image is the frequency below which half of this averaged power lies, and we compared it between V1 and V4 across images with a two-sided sign test. This test treats images as independent observations for fixed encoding and diffusion models, so it does not include variability across animals or model fits; repeating the analysis with a second encoding model (ResNet-50) gave the same result.

#### Comparison of eigenvectors

To compare two geometries at the same image, we measured how closely the subspaces spanned by their leading *k* eigenvectors were aligned. Writing ***V*** _*A*_ and ***V*** _*B*_ for the *P* × *k* matrices whose columns are the leading *k* eigenvectors of the two metric tensors, the subspace alignment is

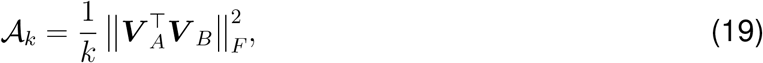

where ∥·∥_*F*_ is the Frobenius norm, the square root of the sum of squared entries. *A*_*k*_ is the mean squared cosine of the principal angles between the two subspaces: it equals 1 when they coincide and 0 when they are orthogonal, and is about *k/P* for random subspaces. It depends only on the subspaces, so it is unaffected by the sign of each eigenvector or by how the eigenvectors are ordered within the subspace.

For Fig. 7E–F, we computed *A*_*k*_ between the two encoding models fitted to the same area, and between the same encoding model fitted to V1 and to V4, separately for the multi-scale and Fisher geometries, and averaged over the 200 test images. We show *k* = 1, …, 50 for the multi-scale (Fig. 7E) and the Fisher geometry (Fig. 7F). To test the difference between the two kinds of comparison, we took for each image the mean alignment of the two same-area comparisons minus that of the two cross-area comparisons, at *k* = 10, and applied a two-sided Wilcoxon signed-rank test across images. As for the spectra, this test treats images as independent observations for fixed encoding and diffusion models.

## Lead contact

Requests for further information and resources should be directed to and will be fulfilled by the lead contact, Matthew Chalk.

## Supplementary note: The multi-scale geometry and mutual information

Here we prove the two relations between the multi-scale geometry and mutual information stated in the main text: the identity 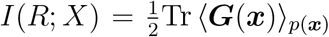, and its geometric form, in which mutual information is proportional to the mean squared magnification of stimulus space. The first proof follows Laquitaine et al.^43^ and the second Azeglio et al.^37^. We restate both in the notation of the main text and make each step explicit.

### Setting and notation

A stimulus ***x*** ∈ ℝ^*n*^ is drawn from a prior *p*(***x***) and evokes a response ***r***, discrete or continuous, with likelihood *p*(***r***| ***x***). We assume only that the stimulus has finite variance, ⟨∥***x***∥^2^⟩ _*p*(***x***)_ *<*∞. Nothing is assumed about the response distribution: the noise may be Gaussian or Poisson, correlated or independent across neurons, and the population may be of any size.

The blurred stimulus is 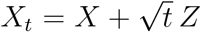, where *Z* ~ *N*(**0, *I***) is independent of both *X* and *R*, so that *p*(***x***_*t*_ | ***x***) = *N*(***x***, *t****I***) as in the main text. Since the blur is added indJependently of the for the blurred prior and *p*_*t*_(***x***_*t*_ | ***r***) = ∫*p*(***x***_*t*_ | ***x***) *p*(***x*** | ***r***) *d****x*** for the blurred posterior. The likelihood response, *R* and *X*_*t*_ are conditionally independent given *X*. We write *p*_*t*_ (***x***_*t*_) = *p*(***x***_*t*_ | ***x***) *p*(***x***) *d****x*** seen by the blurred observer, *p*_*t*_ (***r***| ***x***_*t*_), is defined in Eq. 3 of the main text; by conditional independence it obeys Bayes’ rule in the form

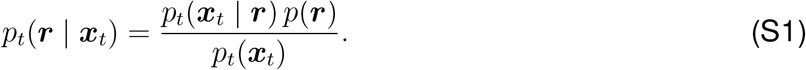

***J*** _*t*_(***x***_*t*_) denotes the Fisher information of *p*_*t*_(***r***| ***x****t*), obtained by applying Eq. 2 of the main text to this distribution.

The proof also uses a second, distinct kind of Fisher information: that of a probability density with respect to translations of its argument,

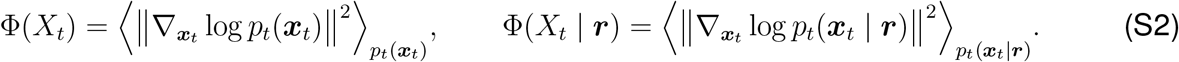

Whereas ***J*** _*t*_ measures how sensitive the responses are to the stimulus, Φ measures how sharply structured a stimulus density is. The central step of the proof relates the two. We use two standard properties of Gaussian blurring, valid for any random vector with finite variance and any *t >* 0^96,97^:

*Fact 1, de Bruijn’s identity*. Differential entropy increases under blurring at a rate set by 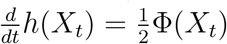 Equivalently, for 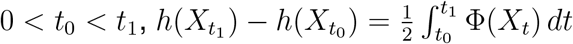, with *h*(*X*_t_) finite for all *t >* 0.

*Fact 2*. Adding independent Gaussian noise of variance *t* leaves at most the Fisher information of the noise itself: Φ(*X*_*t*_) ≤ *n/t*.

#### S1. Mutual information equals half the mean trace of the multi-scale metric

##### Proposition S1.

Under the assumptions above, for every *t >* 0,

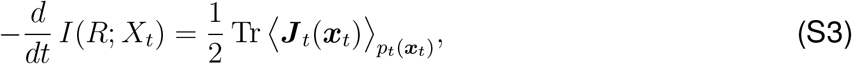

and

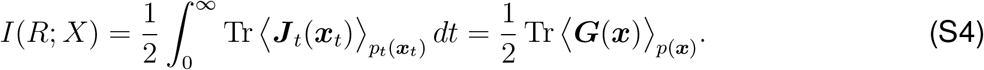

##### Step 1: de Bruijn’s identity for the prior and the posterior

Because *Z* is independent of *X* and *R*, conditioning on *R* = ***r*** leaves the blur unchanged: given ***r***, *X*_*t*_ is a sample from the posterior *p*(***x*** | ***r***) plus Gaussian noise of variance *t*. The blurred poster ior is therefo re the posterior, blurred. The posterior has finite variance for almost every ***r***, since 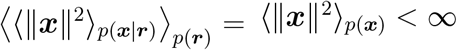. Fact 1 thus applies to both *p*_*t*_(***x***_*t*_) and *p*_*t*_ (***x***_*t*_ | ***r***): for 0 *< t*0 *< t*_1_,

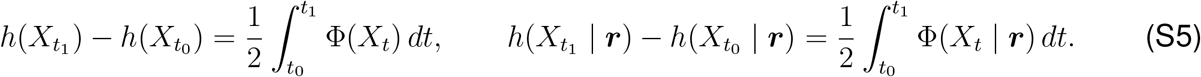

##### Step 2: the rate at which blurring destroys information

For *t >* 0 all entropies are finite, and *I*(*R*; *Xt*) = *h*(*X*_*t*_) − ⟨*h*(*X*_*t*_ | ***r***)⟩_*p*(***r***)_. Averaging the second identity in Eq. S5 over *p*(***r***), which is permitted because Φ ≥ 0, and subtracting it from the first,

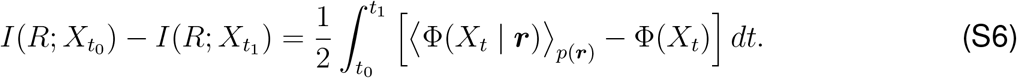

By Fact 2 both terms in the bracket are at most *n/t*, so the integral is finite.

##### Step 3: from the structure of stimulus densities to neural sensitivity

We now show that, for every *t >* 0,

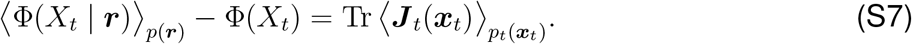

Taking the logarithm of Eq. S1 and its gradient with respect to ***x****t*, noting that *p*(***r***) does not depend on ***x****t*,

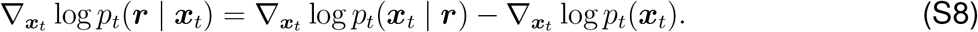

Next, differentiating *pt*(***x****t*) = ∫*p*_*t*_ (***x***_*t*_ | ***r***) *p*(***r***) *d****r*** under the integral, which is justified for *t >* 0 because each *p*_*t*_ (***x***_*t*_ | ***r***) is a Gaussian convolution, and dividing by *pt*(***x****t*),

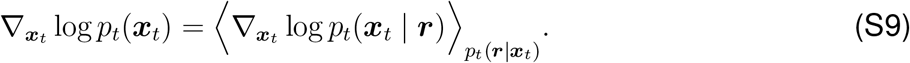

The score of the blurred prior is thus the average score of the blurred posteriors, and by Eq. S8 the score of the likelihood is the deviation of the posterior score from this average. The Fisher information ***J*** _*t*_(***x****t*), the covariance of the likelihood score over *p*_*t*_(***r*** | ***x***_*t*_), is therefore the covariance of the posterior score,

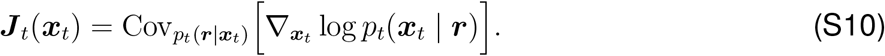

The trace of a covariance is the mean squared norm minus the squared norm of the mean, so

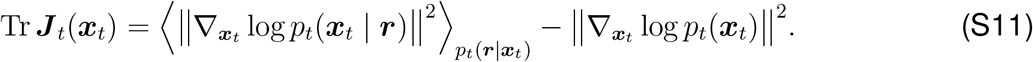

Averaging over *p*_*t*_(***x***_*t*_), and using *p*_*t*_(***x***_*t*_) *p*_*t*_(***r*** | ***x***_*t*_) = *p*(***r***) *p*_*t*_(***x***_*t*_ | ***r***) in the first term, gives Eq. S7.

##### Step 4: integrating over all scales

Substituting Eq. S7 into Eq. S6,

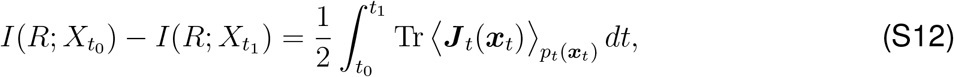

and differentiating with respect to *t*1 gives Eq. S3. It remains to take the two limits. As *t*1→∞, the data processing inequality for the chain 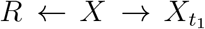, together with the capacity of the Gaussian channel^97^, gives

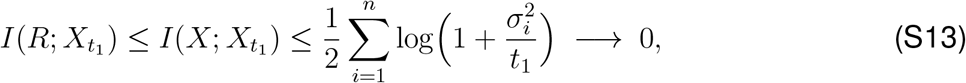

where 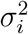 is the variance of the *i*-th stimulus component. As 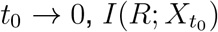 increases, since a stimulus blurred at scale *t*′ *> t* is the stimulus blurred at scale *t* and then blurred further, and data processing applies. It is bounded above by *I*(*R*; *X*), and, since 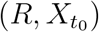 converges in distribution to (*R, X*), lower semicontinuity of mutual information^98^ gives 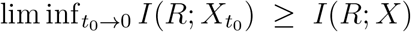. Hence 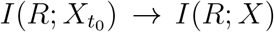. The integrand in Eq. S12 is the trace of a positive semi-definite matrix and so non-negative, and monotone convergence yields the first equality in Eq. S4.

##### Step 5: from the blurred prior to the multi-scale metric

Since *pt*(***x*** _*t*_) =∫*p*(***x*** _*t*_ | ***x***) *p*(***x***) *d****x***, the average over the blurred prior can be taken in two stages, 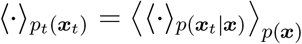. The integrand is non-negative, so the integral over scales and the average over *p*(***x***) can be exchanged, giving

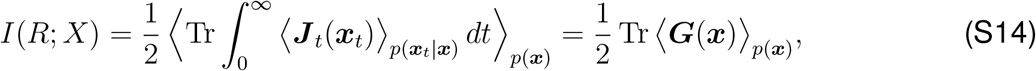

by the definition of ***G*** in Eq. 4 of the main text. ■

#### S2. Mutual information is the mean squared magnification of stimulus space

We now express Eq. S4 in terms of distances. For two stimuli, the magnification factor is *m* = *d****G***(***x, x***′)*/*∥***x*** − ***x***∥′, the ratio of their distance in the multi-scale geometry to their Euclidean distance. We assume that ***G*** is continuous, so that geodesic distances between nearby stimuli are given by the line element of Eq. 1 in the main text,

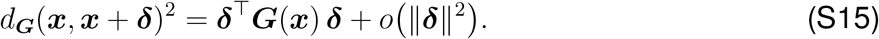

##### Proposition S2.

(a) For small random perturbations 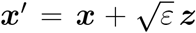, with ***x*** ~ *p*(***x***) and ***z*** ~ *N* (**0, *I***),

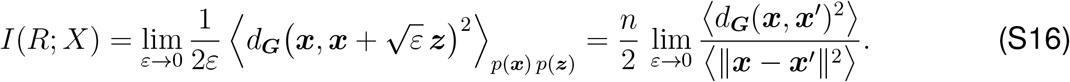

(b) If, in addition, *p*(***x***) has a continuous density on ℝ^*n*^, then for *N* stimuli ***x***1, …, ***x***_*N*_ drawn independently from *p*(***x***),

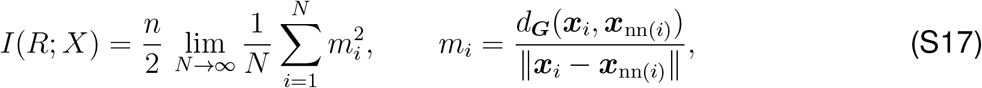

where ***x***nn_(*i*)_ is the Euclidean nearest neighbour of ***x****i* among the samples. That is, 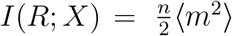 in the limit of dense sampling.

**Proof of (a)**. By Eq. S15, 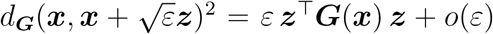. Since ***z*** is independent of ***x*** and ⟨***zz***^⊤^⟩ = ***I***,

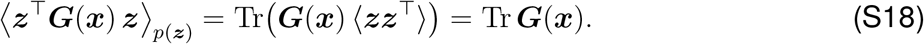

Dividing by 2*ε*, averaging over *p*(***x***) and letting *ε* → 0 gives 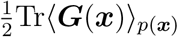, which equals *I*(*R*; *X*) by Proposition S1. The second form follows from ⟨∥***x*** − ***x***′∥^2^⟩ = *ε*⟨∥***z***∥^2^⟩ = *nε*. ■

**Proof of (b)**. Write the displacement to the nearest neighbour as ***x***nn_(*i*)_ − ***x***_*i*_ = *ρ*_*i*_***u***_*i*_, with *ρ*_*i*_ *>* 0 and ∥***u***_*i*_∥ = 1. As *N* → ∞, *ρ*_*i*_ → 0 wherever *p*(***x***_*i*_) *>* 0, so by Eq. S15

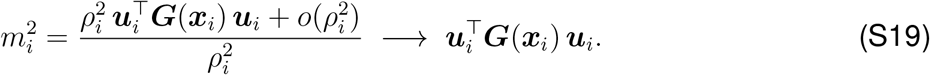

In a small neighbourhood of ***x****i*, the other samples converge to a homogeneous Poisson process with intensity *Np*(***x***_*i*_). Such a process looks the same in every direction, so the direction to the nearest neighbour is uniformly distributed on the unit sphere, independently of ***x****i*, and ⟨***uu***^⊤^⟩ = ***I****/n*. Averaging over samples,

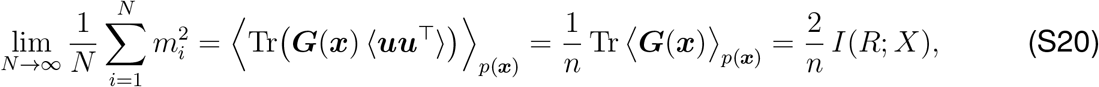

using Proposition S1 in the last step. ■

#### S3. Estimating the multi-scale geometry with diffusion models

For high-dimensional stimuli such as natural images, the integral over stimuli that defines *p*_*t*_(***r*** | ***x***_*t*_), Eq. 3 of the main text, is intractable, and so is direct evaluation of ***J*** _*t*_. Following Azeglio et al.^37^, we instead derive an estimator of the metric in terms of posterior means, which a conditional diffusion model learns.

Writing the average over *p*(***x***_*t*_ ***x***) in Eq. 4 of the main text as an average over Gaussian noise, the squared line element along any stimulus direction ***dx*** is

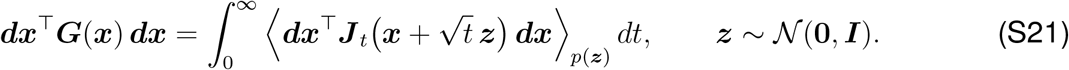

The integral over scales is evaluated numerically on a grid of scales, and the average over ***z*** by Monte Carlo sampling.

##### Neural sensitivity from posterior means

Eq. S10 expresses ***J*** _*t*_(***x***_*t*_) as the covariance, over responses, of the posterior score 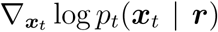. Under Gaussian blur, Tweedie’s formula^90^ relates this score to the posterior mean,

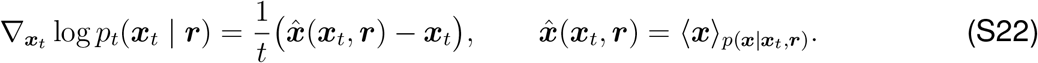

Since ***x****t* does not depend on ***r***, substituting into Eq. S10 gives

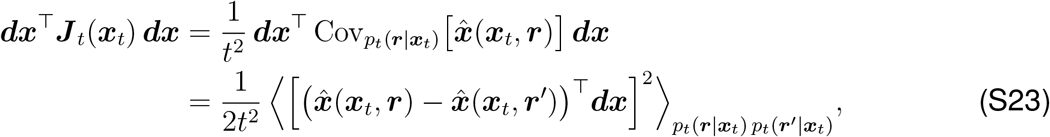

where ***r*** and ***r***′ are two independent responses to the same blurred stimulus, and the second line uses that a variance is half the mean squared difference between two independent draws. The population is thus sensitive to a stimulus direction at scale *t* when two responses to the same blurred stimulus lead to different estimates of the stimulus along that direction.

##### Learning the posterior mean

The posterior mean 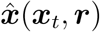 is exactly what a conditional diffusion model learns when trained to denoise stimuli given neural responses^55^. Because Eq. S23 depends only on differences between two outputs of the same conditional model, evaluated at the same blurred stimulus, systematic errors common to both cancel. This avoids the bias of estimators that subtract the outputs of separately trained conditional and unconditional models, which can become unstable as *t* → 0^43,83,99^.

##### Implementation with a standard diffusion model

Denoising diffusion models are usually trained with the variance-preserving parametrization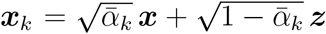, where *k* indexes the diffusion step and 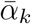 is set by the noise schedule^55^. Rescaling gives 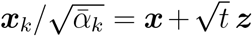 with

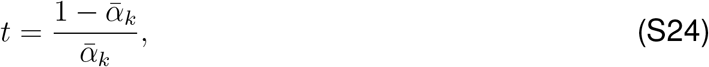

so that each diffusion step corresponds to a blur scale. Since ***x****k* and ***x****t* differ only by an invertible rescaling, conditioning on either gives the same posterior. A network 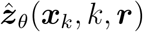, *k*, ***r***) trained to predict the noise ***z*** by minimising the squared error returns, at the optimum, 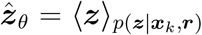, so that

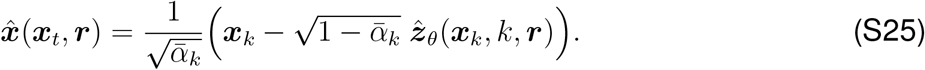

Differences between posterior means are then differences between noise predictions, 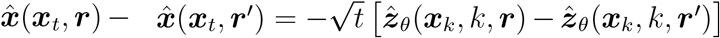. Substituting into Eq. S23, and changing the integration variable in Eq. S21 to log *t*, with *dt* = *t d* log *t*,

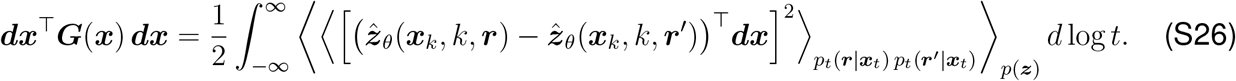

All factors of *t* cancel, leaving squared differences between noise predictions. If the response is withheld on a random fraction of training steps, the same network also returns the unconditional prediction, and hence 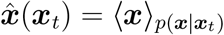^95^.

##### Sampling responses

Eq. S26 requires responses drawn from *p*_*t*_(***r*** | ***x****t*). Exact sampling would first draw a stimulus from *p*(***x*** | ***x***_*t*_), by running the full generative process of the diffusion model, and then a response from *p*(***r*** | ***x***), which is prohibitively expensive. Following previous work^43,100^, we instead use 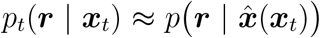, where 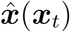 is obtained in a single forward pass of the network. This approximation becomes exact as *t* → 0, where *p*(***x*** |***x***_*t*_) concentrates around its mean. At coarse scales, where it is least accurate, contributions to the metric are small, since *p*_*t*_(***r*** | ***x***_*t*_) approaches the marginal response distribution *p*(***r***) whatever the blurred stimulus.

Each Monte Carlo sample of ***z, r*** and ***r***′ at a given scale contributes one rank-one term to Eq. S26. The estimate therefore takes the form ***G***(***x***) ≈ ***D***^⊤^***D***, and its leading eigenvectors follow from the singular value decomposition of ***D***, without forming ***G***(***x***) explicitly, as described in the Methods.

